# The Roles of Space and Stochasticity in Computational Simulations of Cellular Biochemistry: Quantitative Analysis and Qualitative Insights

**DOI:** 10.1101/2020.07.02.185595

**Authors:** M. E. Johnson, A. Chen, J. R. Faeder, P. Henning, I. I. Moraru, M. Meier-Schellersheim, R. F. Murphy, T. Prüstel, J. A. Theriot, A. M. Uhrmacher

## Abstract

Most of the fascinating phenomena studied in cell biology emerge from interactions among highly organized multi-molecular structures and rapidly propagating molecular signals embedded into complex and frequently dynamic cellular morphologies. For the exploration of such systems, computational simulation has proved to be an invaluable tool, and many researchers in this field have developed sophisticated computational models for application to specific cell biological questions. However it is often difficult to reconcile conflicting computational results that use different simulation approaches (for example partial differential equations versus particle-based stochastic methods) to describe the same phenomenon. Moreover, the details of the computational implementation of any particular algorithm may give rise to quantitatively or even qualitatively different results for the same set of starting assumptions and parameters. In an effort to address this issue systematically, we have defined a series of computational test cases ranging from very simple (bimolecular binding in solution) to moderately complex (spatial and temporal oscillations generated by proteins binding to membranes) that represent building blocks for comprehensive three-dimensional models of cellular function. Having used two or more distinct computational approaches to solve each of these test cases with consistent parameter sets, we generally find modest but measurable differences in the solutions of the same problem, and a few cases where significant deviations arise. We discuss the strengths and limitations of commonly used computational approaches for exploring cell biological questions and provide a framework for decision-making by researchers wishing to develop new models for cell biology. As computational power and speed continue to increase at a remarkable rate, the dream of a fully comprehensive computational model of a living cell may be drawing closer to reality, but our analysis demonstrates that it will be crucial to evaluate the accuracy of such models critically and systematically.

## INTRODUCTION

21st-century cell biology has been transformed by rapid development of new technologies that have delivered our field into an era where scientists can now easily generate vast amounts of quantitative data, providing a broad and comprehensive view of problems that were previously accessible only by brief, laboriously achieved glances through tiny chinks. Northern blots have been replaced by RNA sequencing, Western blots are being replaced by proteomics, and the central cell biological tool of imaging has been revolutionized by a wealth of dramatic improvements in labeling methods, optical design, and digital imaging. While currently available data are just a tiny fraction of the amount that will eventually be needed to model entire cells, sufficient data is available for a number of cell biological processes to ask whether and how we can use it to understand the biological mechanisms underlying those processes.

Mathematical modeling can help (Cohen, 2004; Gunawardena, 2014). By generating a model of a process that we want to study, where we are able to define and control all the inputs and parameters, we can directly determine whether a mechanism we have hypothesized is actually sufficient to explain the phenomenon we have observed. While mathematical modeling of cell-biological processes can take many forms, here we will focus exclusively on computational simulation. One of our goals is to discuss the current state-of-the-art in computational cell biology and remaining open challenges, particularly regarding the spatial organization of signaling processes, the inclusion of stochastic effects, and the multi-scale nature of many cell-biological processes.

Computers are much better than humans at keeping track of large, complex systems with many interacting parts, and also at mercilessly following predetermined rules. One sub-field of cell biology that has been able to make great use of these abilities is the study of signal transduction, where typically many different molecular species interact with one another in branching and reticular networks. Biochemical and genetic experiments have been able to map out and characterize many individual pairwise interactions in multicomponent signaling pathways. However, the resulting “wiring diagrams” have provided little direct insight into how these networks of molecular interactions could give rise to the diverse and fascinating outputs of these systems, which might be able to generate oscillations, high-pass or low-pass filtering of receptor-mediated input signals, conversion of analog signals into switch-like binary outputs, etc. Computational simulations have been instrumental in bringing order and insight into this tangled web so that now it is possible to recognize recurrent motifs in the design of signal transduction systems, and, sometimes, accurately predict cellular responses to external stimuli (Cao et al., 2016; Eungdamrong and Iyengar, 2004; Janes and Lauffenburger, 2013; Kestler et al., 2008).

Within this context, a very fertile ecosystem of computational tools for modeling cellular biochemistry has flourished (Bartocci and Lio, 2016; Gillespie, 1977a).There are now also approaches that enable scientists who are not computational experts themselves to translate their hypotheses about cellular signaling mechanisms into formal models in compact and intuitive ways; these include using textual rules (Faeder, 2011; Harris et al., 2016; Maus et al., 2011) or iconographic symbols (Schaff et al., 2016; Sekar et al., 2017; Zhang et al., 2013) to specify molecular interactions. Based on those specifications, the tools generate the resulting computational representations of the signaling networks and allow modelers to easily modify their assumptions to explore the consequences of such simulated manipulations on cellular behavior. To allow for tool-independent formulation and sharing of computational models, the widely-used “systems biology markup language” SBML was developed. SBML is continuously evolving and is supported by a large number of software systems for simulation and data analysis (Gillespie, 1976a; Gillespie, 1976b; Gillespie, 1977a; Gillespie, 1977b; SBML.org, 2020). While there are many variations, generally simulations of this kind are able to keep track of concentrations and interactions of many individual molecular species as they change over time, often the most interesting dimension for the study of signal transduction.

However, many cell biological processes, including some kinds of signaling, cannot be analyzed without the notion of space. Cell-cell communication is frequently based on the exchange of soluble messenger molecules, such as hormones, cytokines or chemokines, that diffuse through extracellular space before being captured by specific receptors at particular locations on cellular membranes. Direct cell-cell contacts also typically involve only a few of the receptors on a cell and generate localized signals that activate cascades of protein interactions and modifications to propagate from the membrane into the cytoplasm. Spatial simulation of cell biological phenomena is not new; in 1952, Hodgkin and Huxley simulated the propagation of an action potential down a neuronal axon (Gillespie, 1977a; Hodgkin and Huxley, 1952), and in the same year Alan Turing used computational simulation to demonstrate how chemical systems featuring both diffusion and reaction could generate regular spatial patterns from an initial uniform state (Turing, 1952). More recently, many researchers have developed computational simulations that employ state-of-the-art knowledge about the properties and interactions of individual molecular components to attempt spatially-resolved reproduction of complex cell biological phenomena. Reaction-diffusion models of intracellular biochemistry have been used to explore a variety of cellular symmetry-breaking processes, including the establishment of cell polarity (Jilkine and Edelstein-Keshet, 2011). Although the exact mechanistic details vary, spatial simulation approaches have yielded insights for symmetry-breaking systems as diverse as yeast bud site selection (Wedlich-Soldner et al., 2003), asymmetric cell division in early *C. elegans* embryos (Dawes and Munro, 2011) and neutrophil chemotaxis (Onsum and Rao, 2007). Establishment of spatial gradients that determine cell fate has been explored in cells ranging from giant syncytial *Drosophila* embryos (Gregor et al., 2007) to tiny individual bacteria (Chen et al., 2011). Simulations that explicitly consider spatial effects as one cell communicates with its neighbors have been used to understand the formation of regular stripes in *Drosophila* embryos (von Dassow et al., 2000), and bizarre non-cell-autonomous effects in the patterning of wing bristles in adult flies (Amonlirdviman et al., 2005).

In many of the modeling efforts that we have mentioned so far, the computational model and/or simulation was formulated explicitly for the problem at hand. While this approach has enabled important scientific insights, we believe that spatial simulation for cell biological processes can become a much more widely used tool in the cell biologists’ toolbox if there were more general access to user-friendly implementations of general spatial modeling frameworks that do not require extensive computational expertise to use. Consider, for example, the wide variety of user-friendly open-source software packages now available for analysis of sequencing data (Rice et al., 2000; Trapnell et al., 2012). One particularly important benefit of more standardized approaches to spatial simulations in cell biology is that standardization may help to resolve whether conflicting conclusions arise because of fundamental scientific differences in model assumptions or because of details of numerical implementation.

Another aspect gaining importance as we zoom in closer on the building blocks of cellular structures is the fact that, at the molecular level, cellular biochemistry is governed by stochastic processes such as thermal Brownian motion and the collisions and interactions among individual particles (Schnoerr et al., 2017). Only in the limit of high concentrations and homogenous spatial distributions can the behavior of the molecular components of cellular signaling pathways be described in terms of deterministic reaction rate equations. Many subcellular mechanisms operate far from this limit either due to highly non-homogenous clustering of receptors and of the signaling components they recruit, such as studied in the MAPK signaling pathway (Takahashi et al., 2010), or due to locally low copy numbers, for instance of multi-molecular complexes regulating transcription in the nucleus (Cho et al., 2018). To accurately capture the stochastic characteristics of such processes, computational models have to simulate the motion and interactions of individual molecules. A full simulation of Brownian dynamics, following each single ‘Brownian hop’ of all molecules of a cellular region would, however, in most cases be too computationally expensive and time-consuming. Moreover, by choosing a particular time step for recreating Brownian hops on the computer, we would impose this time scale on our simulations, missing events, such as molecular encounters, that may occur ‘in between’ our time steps. Many approaches have been developed to deal with this problem and we will discuss several of them below. A common theme among all of them is, however, saving computational cost through temporal and spatial coarse graining without sacrificing too much accuracy in the simulation results. This challenge unites researchers looking at spatially-resolved modeling (without a focus on stochastic effects) and those that try to capture the manifestations of stochastic fluctuations in cellular systems.

The question of how the field could best go about building and sharing broadly applicable computational tools for spatial and stochastic modeling of cell biological processes was the focus of our working group, organized by J.R.F. and R.F.M., which met with the support of the National Institute for Mathematical and Biological Synthesis (NIMBioS). We chose to focus specifically on the biochemical scale of molecular interactions. We did not attempt to include the enormous field of molecular dynamics (MD) simulations that explores forces and movements of individual atoms within proteins or other macromolecules. The reason is that, because of their intensive computational demands, MD simulations are currently limited to exploration of very small biological systems (a few macromolecules) over very short periods of time (typically in the sub-microsecond range), too small and too fast to be incorporated into cell-scale computational simulations. Conversely, we also limited ourselves to considering simulations of interactions within systems of molecular complexes with specific stoichiometry rather than extending our analysis to mesoscopic-scale models that abstract the behaviors of complex molecular systems into continuum physical descriptions, such as those describing cytoskeletal filaments as elastic beams (Nedelec and Foethke, 2007a; Odell and Foe, 2008) or the plasma membrane as a flexible thin film (Fowler et al., 2016). While these kinds of models are enormously useful in cell biology, they rely on fundamental simplifying assumptions. In contrast, we are specifically interested in exploring whether detailed simulations of the behaviors and interactions among biσ molecular complexes can succeed in predicting certain kinds of mesoscopic phenomena, and hence may help determining under which conditions the simplifying assumptions are justified. As we will discuss in our conclusion, it will be an exciting future direction for the field of biological simulation when all these three levels of spatially-resolved simulations can be seamlessly interconnected.

In this article, we will first briefly survey several existing approaches for spatial and stochastic cell simulations and then apply them to a series of ‘unit tests’ and benchmark problems. The problems we chose are categorized to cover a variety of different aspects of spatially resolved and/or stochastic simulations of cellular behavior. Conceptionally, they are relatively simple and are meant to capture specific challenges related to essential aspects of biological processes. They must be accurately modeled by a simulation tool to ensure that that tool’s results are reproducible for the problem category covered by the unit tests or benchmark problems.

With these examples in hand, we then summarize how features of a particular cell biological problem should guide selection of the appropriate modeling approach. Finally, we will present the results of our wide-ranging and, sometimes, highly opinionated discussions on future directions and challenges for the field. We all share an ambitious vision of the future power of spatial cell simulations both for exploring hypotheses about mechanisms and for coming to grips with the massive amounts of quantitative data now available to cell biologists. Our overall goal here is to map out where the field currently stands and propose a trajectory for the future.

## MODELING APPROACHES

### Overview of approaches for spatial and stochastic modeling and simulation of molecular reactions underlying cell-biological phenomena

Computational models that simulate the biochemistry underlying cell-biological processes need to be able to describe molecular players and their reactions. However, depending on the particular question at hand, taking into account spatial aspects and stochastic effects (Fig 1) may or may not be essential, as we show in the Results section. The addition of spatial resolution is computationally demanding, and stochastic simulations are usually more costly than their deterministic counterparts. To attempt a whole-cell simulation, for example, one must choose whether more components and a more complex reaction network is necessary, generally requiring the sacrifice of spatial resolution (Sanghvi et al., 2013) or if spatial resolution is necessary, then the reaction network must be simplified (Ghaemi et al., 2020). If spatial resolution is a priority, must species be resolved as individual particles, capturing fluctuations in copy numbers but at considerable extra expense (Table S3), or is an efficient deterministic approach sufficient?

**Fig. 1:**
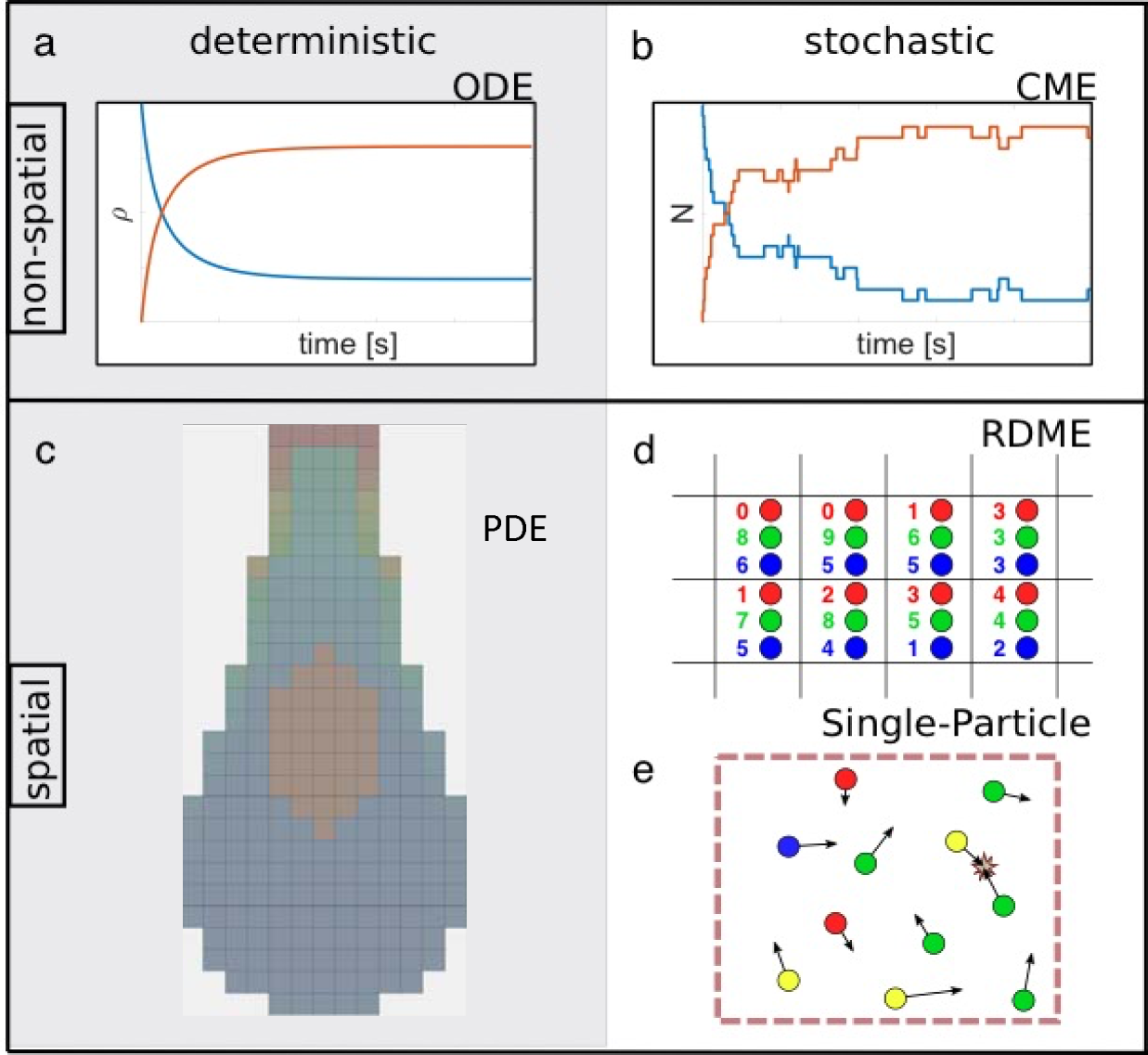
Overview of non-spatial and spatial simulation approaches for describing the time-dependence of interacting and reacting species. Different colors represent different species. Spatial models are illustrated at one point in time. Left: Deterministic approaches to modeling biological systems solve a) ordinary or c) partial differential equations (ODE/PDE) using standard numerical methods, benefiting from extensive method development across science, math and engineering fields. PDEs are numerically solved on a mesh as shown. Right: Stochastic simulation approaches sample from a time-(and space-) dependent probability distribution that typically models unimolecular and bimolecular reactions, as well as diffusion for spatial methods. d) Reaction-Diffusion Master Equation (RDME) methods are the extension of the b) Chemical Master Equation (CME) methods on to a spatial lattice, where integer copy numbers of species are tracked and can diffuse between lattice subvolumes. e) Single-particle methods propagate individual particles undergoing diffusion in continuous space, where bimolecular reactions can occur only upon collision or co-localization in space. We provide a guide to corresponding software tools of each approach in Table S1.

### Non-spatial modeling approaches

In many situations, we can describe a biochemical system adequately in terms of the overall concentrations of interacting molecule types and complexes (collectively called ‘species’), while neglecting the spatial variations in these concentrations. Reaction rate equations (**see Box 1**) describe how the species concentrations evolve in time. The terms in these equations arise from the rates of the reactions that can occur in the system, which are often described by the Law of Mass Action. The rate of reaction between two interacting species can be given by the product of their concentrations and a rate constant, e.g., *k*_on_ for the ligand-receptor binding described in Box 1. The bimolecular rate constants that appear in these equations are sometimes referred to as ‘macroscopic’ rate constants because they describe the average rate of reaction assuming homogenous distribution of the reacting species. In contrast, ‘microscopic’ rate constants, which govern reaction kinetics at the scale of interacting particles, may take into account more details about the way the molecules approach each other, as discussed below.

#### Box 1

**Reaction Rate Equations:**

Technically speaking, reaction rate equations are ordinary differential equations (ODEs). Here, the ‘ordinary’ refers to the fact that they involve only time (as opposed to, for instance, time *and* space). To describe the time evolution of multiple interacting molecule types, one uses coupled differential equations that express how the components’ concentration changes are linked (or coupled). For applications, see, for example (Aldridge et al., 2006; Tyson et al., 2003). From a numerical/mathematical point of view, ODEs describing biochemical reactions are typically simple and many tools exist that can solve them to obtain the temporal evolution of the concentrations in ODE models.

Consider a simple model of a receptor binding to a ligand. We call *R* the concentration of the receptor, *L* that of the ligand, and *RL* that of the complex formed by the binding of the two. The rate equations giving the time derivatives of *RL, R* and *L* for this reaction could be written as

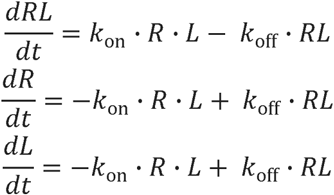

Here, *k*_on_ and *k*_off_ are the association and dissociation constants, respectively. The time course of *RL* would look similar to the red curve in ***Fig***.***1a***, whereas time courses of *R* and *L* would be similar to the blue curve. These equations can be solved analytically, but the additional complexity of most biologically-relevant models generates equations that require numerical solution by computer.

The chemical master equation considers the discrete and stochastic nature of the biochemical system. The CME describes how the probability of the system being in a specific state evolves over time, by using reaction probabilities (likelihood of occurrence per unit time) rather than the equivalent reaction rates. Just like reaction rate equations, the CME assumes well-stirred (homogeneous) systems. In most practical modeling applications the CME cannot be solved analytically, but simulations of the CME are conceptually straightforward and very widely used (**see Box 2**).

#### Box 2

**The Chemical Master Equation (CME):**

The chemical master equation (CME) is a set of coupled linear ODEs that describe the time-dependence of the probability to occupy each of a set of discrete states, with each state defined by the copy numbers of molecular species. The CME may be written as

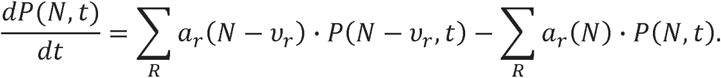

where the composition vector *N* is composed of copy numbers of each molecular species, and thus one has a set of equations, one for each possible instantiation of *N*. Here, the sum runs over all possible reactions. ν_*r*_ is the stoichiometric vector of reaction *r* that describes how this reaction changes the number of molecules in the composition vector *N* and *a*_*r*_ (*N*) is the probability per unit time that reaction *r* occurs, given that the system is in the state described by *N*. The CME is important from a conceptual point of view as it represents a framework to describe probabilistic transitions and thus captures the stochasticity underlying all molecular interactions(Schnoerr et al., 2017) (Grima and Schnell, 2008). The computational cost of solving the CME equations scales exponentially with the number of chemical species, and, although clever approaches have extended the size of systems for which the CME can be solved (Munsky and Khammash, 2006), high computational cost still limits biological applications. An intuitively simple way to calculate a solution of the CME would be to set up a simulation where time ticks forward in small, discrete intervals (time steps). However, the fixed time step would always introduce the potential for error since reactions may occur even during shorter time steps than the one chosen to propagate the system in time.

One popular and precise method used to generate trajectories through the state space sampled by the CME without the need to choose a discrete time step is the Gillespie algorithm, also known as the stochastic simulation algorithm (SSA) (Bortz et al., 1975) (Gillespie, 1976a). In a Gillespie simulation, the time interval until the next reaction occurs is itself sampled, as is the type of reaction that will occur (Gillespie, 1976a). Molecular species that can react quickly and have many possible interaction partners will be selected frequently while rarer molecules associated with slower reactions will be selected rarely. Since the simulations proceed with one reaction at a time, the computational cost depends strongly on the number of particles and reaction rates in the system. In contrast, the effort required to integrate (or solve) reaction rate equations depends mostly on how many molecule types are involved and whether their interactions occur on different or similar timescales. Various approaches have been developed to increase the efficiency of both exact (Gupta and Mendes, 2018) and approximate (Schnoerr et al., 2017) stochastic simulations of the CME. In addition, efficient methods have been developed to compute distributions and moments directly from the CME itself (Hasenauer et al., 2014). A stochastic simulation of the example system from Box 1 would look similar to ***Fig. 1b***. Note, however, that such a trajectory represents only one possible time course compatible with the underlying CME. This means that many stochastic simulation trajectories must be collected to determine probability distributions and moments of the CME.

Both the reaction rate approach and the CME approach simply cannot capture effects of inhomogeneous distributions of molecules in space, such as receptors clustered on membranes, or intrinsic time delays due to diffusion to localized targets such as membrane bound receptors. For more realistic simulations of cellular process, we must turn to different computational approaches that explicitly include space.

### Spatial modeling approaches

The molecular components of living systems are not distributed homogeneously and the high spatial resolution of today’s fluorescence microscopy is continuously giving us more examples of biological phenomena where the spatial arrangement of the underlying biochemical processes is fundamentally important. To model such phenomena we have to switch from non-spatial to spatial simulations. However, this switch is frequently not easy due to a growth in the number of model and system features that must be specified (Fig. 2a). The most important difference between non-spatial and spatial simulations is that the latter take into account the translocation of the interacting molecules. In the simplest case, this means that, in addition to reactions, the diffusion of molecular species in space must be simulated. Along with diffusion, the system geometry must be specified and what happens at the ‘walls’. This is significantly more challenging to implement when the spatial system containing the interacting molecules is not simply a square box with rigid walls. Realistic spatially resolved models often aim to capture aspects such as particular cell morphologies; examples include synaptic structures with narrow regions connected to larger cell bodies (Cugno et al., 2019; Rangamani et al., 2016) or geometries that are flat and almost twσdimensional, like lamellipodial extensions in migrating eukaryotic cells (Nickaeen et al., 2017) (Fig. 2b). Geometries for cell simulations can be designed by hand, derived from microscope images (Schaff et al., 2000), or generated from machine-learned cell models (Majarian et al., 2019). We note that spatial models have the capacity to build in many additional features such as mechanics, electrostatics or coarse molecular structure that may be particularly important for useful simulations of some cell biological processes (Fig. 2d). There are very few existing computational tools that take these additional features into account, largely because they introduce further challenges to both the numerical implementation and the mathematical descriptions of the physical model.

**FIGURE 2:**
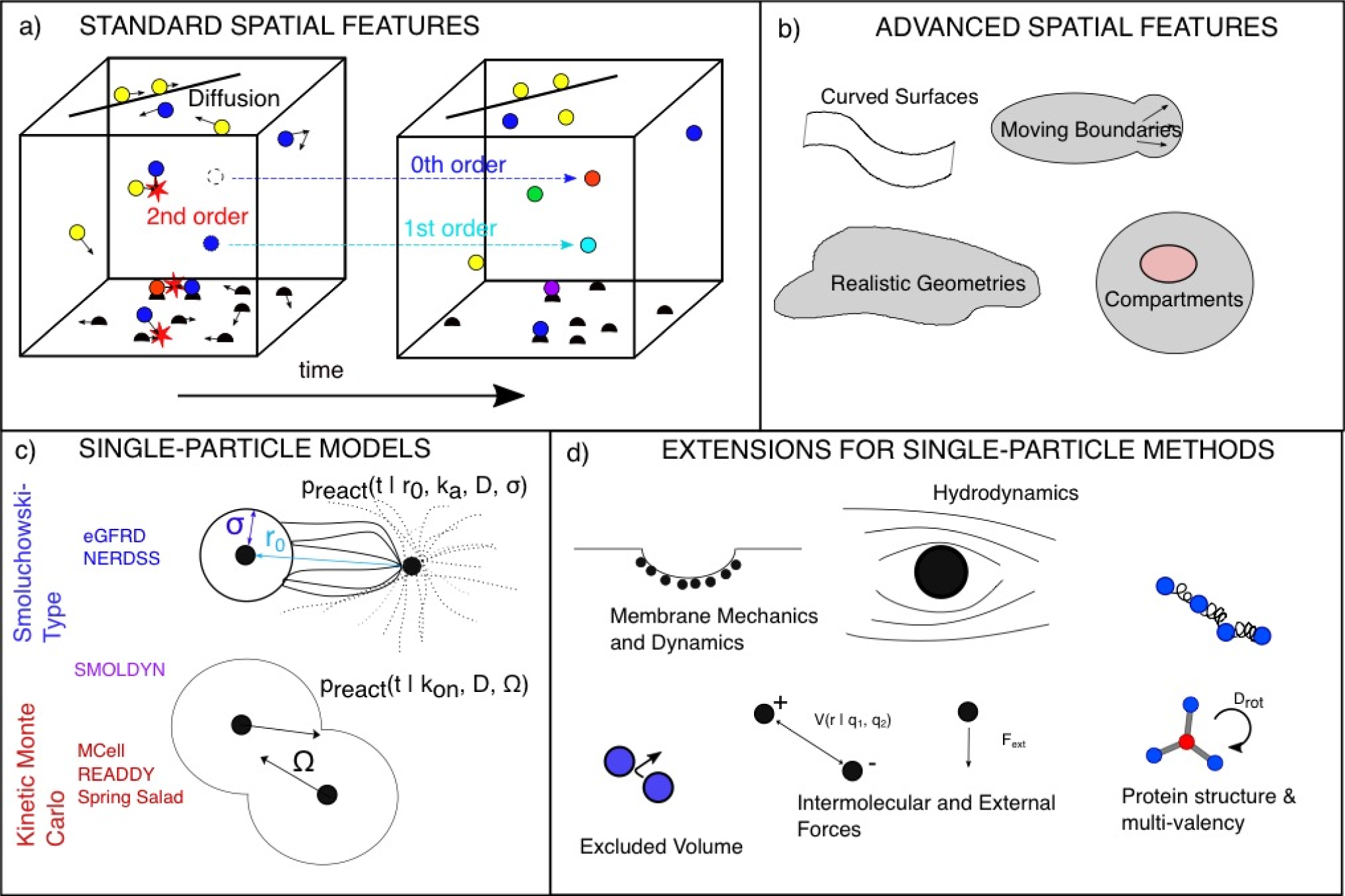
Demands on spatial approaches and their extensibility. a) For all spatial models, in addition to treating fundamental reaction types in all dimensions (3D, 2D,1D) and between all dimensions (for instance molecular exchange between bulk (3D) and surfaces (2D) they must specify an equation of motion, typically diffusion, and boundary conditions on the system geometry. b) More advanced treatments of curved and complex boundaries require additional care for treating reactions and diffusion. c) Single-particle spatial methods fall into two broad classes. In Smoluchowski-type methods, reaction probabilities (p_react_) are determined by finding analytical solutions for the fraction of diffusive trajectories that produce collisions that lead to reactions (not all collisions lead to reactions when the microscopic rate k_a_<∞). In single-particle Kinetic Monte Carlo methods, algorithmic variations exist but particles that are in a neighborhood *Ω* can react according to a macroscopic rate, k_on_. Smoldyn, for instance, is an approximation to the Smoluchowski model. d) Single-particle methods have the capacity to build in higher-resolution features, although these will alter their equation of motion, requiring new definitions of p_react_. We note that PDE-based models can also expand beyond purely diffusive dynamics. In Table S2 we summarize features available in commonly used software tools.

#### Reaction-diffusion equations

represent the most straightforward extension of reaction rate equations for the inclusion of spatial aspects. Instead of depending just on time as a variable, the behavior of molecular species now additionally depends on spatial coordinates. Equations describing these reactions must therefore be formulated as partial differential equations (PDEs) rather than as ordinary differential equations (ODEs) (Fig 1c). As is frequently the case for mathematical models of biological phenomena, only very simple situations can be described through equations that can be explicitly solved in such a way that the solution describes the behavior of the modeled system as a continuous function of space and time (Lipkow and Odde, 2008). In most cases, one has to explore reaction-diffusion equations through numerical simulations that divide space into sub-volumes (frequently called ‘voxels’) and calculate how diffusion leads to exchange among the sub-volumes. Reaction-diffusion equations have been widely used to model spatially-resolved biomolecular dynamics and interactions of cell-biological systems (Loew and Schaff, 2001). The spatial dynamics can be extended beyond pure diffusion (e.g. to include advection), and reactions can be defined phenomenologically (Hill-type or Michaelis-Menten). Like reaction rate equations, deterministic PDE reaction-diffusion equations do not capture stochastic fluctuations in species numbers, and thus cannot, for example, capture pattern formation driven by a system’s sensitivity to low copy numbers (Howard and Rutenberg, 2003).

One way that stochastic reaction-diffusion equations can be formulated is the spatial extension of the chemical master equation, known as the Reaction Diffusion Master Equation (RDME) (Fig 1d). Instead of only defining how a system switches from one set of numbers of molecules in particular states to another (for instance when a complex in the system decays into two molecules) the RDME includes ‘hops’ from one location to another. Importantly, just like the CME, the RDME describes discrete changes. That means, it requires a spatial discretization into sub-volumes, within each of which well-mixed conditions are assumed to prevail. Diffusion events of molecules are tracked only when they occur between adjacent sub-volumes, not within an individual sub-volume (Fange et al., 2010). Similar to the non-spatial case, it is usually not possible to solve the RDME analytically, and instead it is standard practice to compute solutions by simulating a particular stochastic time evolution of the system with, for instance, a Gillespie approach (see Box 2) that adds diffusional hops to the list of events that can occur. However, care must be taken to choose the right degree of spatial resolution that strikes the appropriate balance between capturing spatial details and avoiding sub-volume sizes that are so small that discretization dilutes the molecules to a point where they essentially do not ‘see’ their potential reaction partners anymore because the molecules are spread out over distinct sub-volumes.(**see Box 3**).

##### Box 3

**Spatial Discretization of Reaction Diffusion Equations:**

To numerically solve reaction-diffusion processes modeled as partial differential equations (PDEs), it can be challenging to choose the appropriate spatial discretization of the modeled biological geometry. A discretization that is too coarse will suppress many spatial details and will represent a poor approximation of the underlying biology. However, keeping track of the contents of many very small voxels will not only be very expensive computationally, but may also lead to situations where the assumptions of mass-action kinetics no longer strictly hold since the fraction of the molecules in the system that populate a single voxel becomes so small that the very concept of an average concentration becomes problematic. Furthermore, the accuracy and efficiency of PDE solvers is not just sensitive to the resolution of the spatial discretization (sometimes called lattice or mesh) but also to the discretization scheme as manifested, for instance, in the shape of the voxels. The practice of designing adaptive meshes, that is combining voxels of different shape and size in one simulation to capture small-scale spatial details where needed while keeping the total number of voxels as low as possible, is a field of active research. The structure of the mesh also has to be adjusted to the numerical method chosen to solve the reaction-diffusion PDEs. Finite volume methods directly simulate diffusional exchange between voxels. In contrast, finite element algorithms optimize the coefficients of interpolation functions at the nodes of the mesh to achieve good approximations of concentration profile resulting from the combination of reactions and diffusion. See, for example (Richmond et al., 2005).

Similar to PDEs, the accuracy and cost of the ***Reaction Diffusion Master Equation*** (RDME) is sensitive to the spatial mesh; this problem is inherent to all spatially discretized simulations. Computational costs grow rapidly as the mesh resolution increases. Importantly, the accuracy of an RDME model does not always increase with a finer mesh. A very small mesh size violates the assumption that species are dilute and their own molecular volume is small relative to the voxel (Erban and Chapman, 2009) (Isaacson and Zhang, 2018; Wolf et al., 2010). For specific non-fundamental reaction types, RDME has an additional limitation in that it does not always converge to the CME solutions in the limit of fast diffusion, as expected (Smith and Grima, 2016). Hence, the RDME may be viewed as a *non-converge*nt approximation of more microscopic spatially continuous models, such as the Smoluchowski model to be discussed below.

#### Particle-based spatial simulation methods

take into account the stochastic motion and interactions of individual molecules in continuous time and space and are thus capable of modeling biochemical processes that involve low copy numbers and strongly heterogeneous molecular spatial distributions (Fig. 1e). These methods have the highest resolution (Fig. 2c) but they come with a high computational cost. Importantly, simulations that treat each particle as an individual also offer the possibility of building in more detailed molecular features (Fig. 2d). Typically, particle-based approaches resolve a bimolecular reaction A+B? C of a pair of molecules as two physically distinct stochastic processes. First, the molecules’ diffusive (Brownian) motion leads to their encounter. Then, the molecules either form a bond with a reaction probability determined by the reaction rate constant and their current separation, or else they diffuse away from each other. Numerical approaches to single-particle reaction-diffusion calculations for biological systems can be usefully grouped into two classes; first, what we will call Smoluchowski-based methods, and second, what we will call single-particle Kinetic Monte Carlo (KMC) schemes (Fig. 2c).

The definition of the reaction probability is the primary challenge and distinguishing feature of different single-particle algorithms. The first class of single-particle methods we will consider, Smoluchowski-based methods, derive reaction probabilities from the Smoluchowski model of diffusion-influenced reactions (Collins and Kimball, 1949; von Smoluchowski, 1917). These methods explicitly recognize that the macroscopic kinetics of association must depend on both diffusion and reaction rate constants, resulting in a distinction between a microscopic and macroscopic rate (**see Box 4**).

##### BOX 4

**Microscopic vs Macroscopic Rates, and the Sensitivity of Strong Binding to Diffusion**

For all non-spatial models, as well as PDE, RDME and KMC-type spatial models, bimolecular association reactions are parameterized by the macroscopic rate constants, k_on_, corresponding to the rates one would measure from a binding experiment in bulk solution. This is because in all these spatial models, species that are localized in a small volume are assumed well-mixed, thus obeying the same mass-action kinetics used in non-spatial models. In the Smoluchowski-based single-particle methods, however, molecular interaction kinetics are split into two steps, as described in the text. This results in a purely diffusive contribution to the bimolecular collision, and then what is effectively a purely energetic contribution defined by a microscopic on-rate k_a_, assumed to occur at a specific binding radius. In 3D, this results in the long-known relationship:

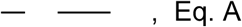

where *D* is the sum of both species diffusion coefficients. With this relationship we can directly assess the impact of diffusion on controlling macroscopic kinetics. As shown in the Figure, for large microscopic rates, the macroscopic kinetics of the A+B→∅ reaction is noticeably dependent on diffusion, whereas for smaller rates, the effect of diffusion is negligible, despite D_A_=D_B_ dropping from 100 to 1 um^2^/s. Here A_0_=B_0_=62 M. Hence large k_a_, or strong binding, is diffusion-limited, and small k_a_ is rate-limited.

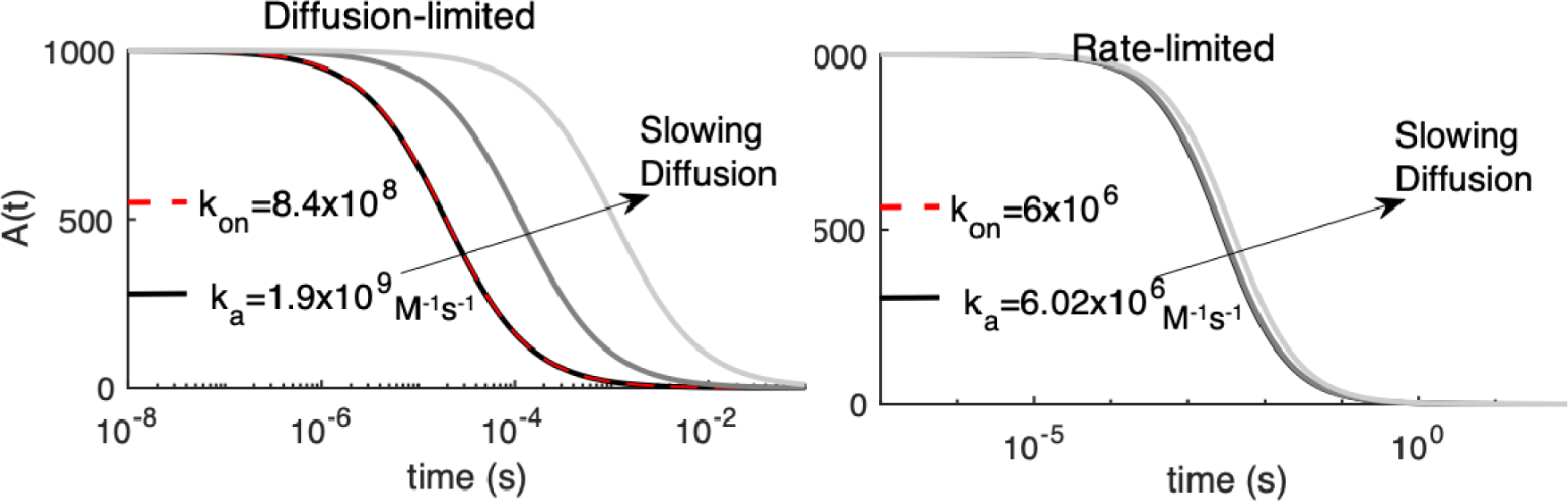

In two and one dimensions (e.g. on surfaces and filaments), the relation between microscopic and macroscopic parameters is more complicated, due to the properties of diffusion in lower dimensions smaller than three. No single relationship exists between microscopic and macroscopic rates [see e.g. (Yogurtcu and Johnson, 2015)], but meaningful theoretical relationships can be defined if the system size is considered (Szabo et al., 1980), where in 2D we further correct for system density using (Yogurtcu and Johnson, 2015):

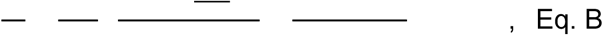

where 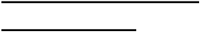.

and are the copy numbers of reactants A,B and the system size is S.

For our test cases below, we thus always derive the microscopic rates k_a_ to reproduce the macroscopic rates using Eq A or Eq B. This is because it is already clear from non-spatial models that changes to the macroscopic rates will necessarily alter the reaction kinetics. Since we are not focused on probing the influence of kinetic parameters on molecular behavior, but rather the role of explicit spatial representations in controlling species distributions and encounter times, we preserve all k_on_ values. However, it is worth noting that for a reaction pair with a large k_a_ value, if diffusion slows throughout the simulation due to, for instance, formation of large complexes, then the macroscopic kinetics will also slow down, an effect that is naturally captured in Smoluchowski-based methods.

Calculating the positions and reactions of all molecules in this framework by “brute force” typically requires computationally costly simulations using extremely small time steps to minimize the risk of missing molecular encounters. An alternative approach uses a mathematical formalism that predicts the encounter probability for pairs of particles based on the particles’ initial positions. This ‘Green’s function’ approach can, however, only calculate the encounter probability for two particles at a time. This means that the simulated system has to be segmented into twσparticle subsystems (**see Box 5**). In practice, this turns out to be feasible for many interesting biological problems. Several computational frrameworks have been developed to take advantage of this approach, including FPR (Johnson and Hummer, 2014), NERDSS (Varga et al., 2020), GFRD (van Zon and ten Wolde, 2005) and eGFRD (Sokolowski et al., 2019) (Fig. 2c). A related computational implementation, known as Smoldyn, approximates the solution to the Smoluchowski model (Andrews, 2017; Andrews et al., 2010; Andrews and Bray, 2004). By approximating the solution, Smoldyn is simpler to implement than the more strictly constrained implementations listed above, but the time-dependence and in some cases equilibrium are not as rigorously correct.

##### Box 5.

**Single-particle reaction-diffusion simulations:**

Single-particle methods simulate the stochastic behavior of individual and (in spite of the name) pairs of interacting molecules. Any biochemical network whose description does not include ad-hoc phenomenological processes (such as, for example, Hill coefficients describing non-linear dose-response characteristics) can be described as composed of uni- and bimolecular reactions. As unimolecular reactions are only time-dependent, they are typically modeled as Poisson processes. For bimolecular reactions, the distance between a pair of particles influences the probability that they will diffuse to collision and react with one another in a time-step. The time evolution of the molecules’ positions is described by a stochastic differential equation, the overdamped Langevin equation(Van Kampen, 2007). Its numerical implementation, known as Brownian Dynamics (BD) (Ermak and Mccammon, 1978; Northrup et al., 1984), requires tiny time steps to accurately resolve molecular encounters, which renders the BD scheme highly inefficient. It is, however, possible to introduce a shortcut by using Green’s function approaches to calculate the distance-dependent reaction probabilities for pairs of molecules. This can enhance the efficiency of BD simulations by resolving bimolecular reactions within one large time step, *Δt*.

For the Smoluchowski model, the Green’s function (GF) *p*(*r,Δt* |*r*_0_) can be obtained as the solution of the diffusion equation that describes a pair of molecules *A,B* that diffuse with diffusion constants *D*_*A*_, *D*_*B*_, respectively, and may undergo a reaction *A + B → C* as follows

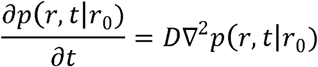

where *D= D*_*A*_ *+ D*_*B*_ and *r, r*_*0*_ refer to the distance between *A* and *s* after and before the time step, respectively. In accordance with the twσstep picture described in the main text, reactions are incorporated by imposing boundary conditions that specify the physics at the encounter distance *r= σ*, such as Smoluchowski-Collins-Kimball BC (Collins and Kimball, 1949)

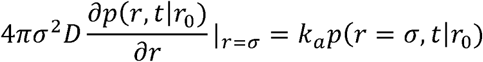

where *k*_*a*_ refers to the intrinsic reaction constant. It is worth noting that finding the appropriate Green’s function to capture the desired properties of the molecular interactions can be a challenge. For the case of *reversible* diffusion-influenced bimolecular reactions of an isolated molecule pair in 2 dimensions the GF was derived only in 2012 (Prustel and Meier-Schellersheim, 2012).

The second class of particle-based approaches we will consider, single-particle Kinetic Monte-Carlo (sp-KMC) schemes, are also based on the twσstep picture of bimolecular reactions. However, the sp-KMC methods do not seek to strictly implement an underlying physical model, unlike the Smoluchowski-based models. Instead, the reaction probabilities are derived by using various approximating assumptions. Widely-used implementations include MCell (Kerr et al., 2008), ReaDDy (Schoneberg and Noe, 2013), and SpringSaLaD (Michalski and Loew, 2016). Within these frameworks, the mathematical expressions of reaction probabilities typically assume a simpler, computationally more efficient form than their Smoluchowski model counterparts. While this is obviously useful for improving computational speed, it may not be immediately clear to what extent the approximations are justified and how much error they introduce. While both classes of model can predict the same behavior in simple systems (Tauber et al., 2005), Smoluchowski-based algorithms inherently capture excluded volume (particles exclude a sphere of radius *σ* relative to partners) and microscopic collisions, whereas sp-KMC methods do not. Smoluchowski models thus provide higher resolution at short distances and for dense systems.

The future extensibility of both classes of approach hinges on the feasibility of finding mathematical expressions for the crucial reaction probabilities that incorporate additional features and details, such as curved surfaces, intra-molecular constraints, and external and internal deterministic forces (Fig 2b and 2d). Arguably, this task is easier for sp-KMC methods, because they rely on approximations rather than exact analytical solutions to derive these expressions. However, as discussed, this flexibility may come at the expense of a clear physical justification of the underlying assumptions and rigorous error control.

## RESULTS

We present a series of test problems relevant to spatial modeling of cellular and subcellular processes. This list is not meant to be exhaustive, but rather to permit a manageably-sized survey of the kinds of problems that different simulation programs may be challenged to solve within a larger biological study. Our selection thus includes very simple problems that can be solved by many simulation tools as well as complex problems that can be solved (at present) by only a few. By framing the problems explicitly, we facilitate a direct comparison among different simulation packages both with respect to accuracy of execution and how they encode these particular scenarios. While all the models presented have been simulated previously, they have not been subject to the quantitative comparative analysis performed here across multiple model and method types. This comparison provides us with distinctive insight into the sensitivity of quantitative and qualitative behavior that emerges with specific biologically relevant features, which we summarize at the end of the Results section.

For these test cases, we contrast results from stochastic, deterministic, spatial and non-spatial modeling approaches. We use Virtual Cell software for all non-spatial simulations and spatial deterministic simulations. For spatial stochastic simulations, we use the single-particle softwares NERDSS, Smoldyn, MCell, and eGFRD. We provide executable model inputs and numerical outputs for each model in our publicly accessible repository (https://github.com/spatialnimbios/testcases/). We summarize a broader range of actively developed tools and their features in Table S2, as distinct software tools have introduced selected complex features of RD systems (also discussed previously (Schoneberg et al., 2014; Takahashi et al., 2005)), and this is an additional consideration for users when selecting a tool for their biological problem.

### Category 1: “Unit test” cases

This category of problems represents fundamental building blocks for which there is a known correct answer (at least at steady-state). We emphasize that because these reactions form the basis of much more complex models and geometries, it is essential that they be carefully tested for accuracy with regard to reaction kinetics and, in the case of bimolecular reactions, reversibility. For all of these, we initialize simulations with well-mixed components, and thus one may expect that any modeling approach would give the same outcome. However, due to differences in both approaches and algorithmic choices, we find that differences do in fact emerge in specific parameter regimes, particularly at short times before the system approaches steady state (Fig. 3).

**Figure 3.**
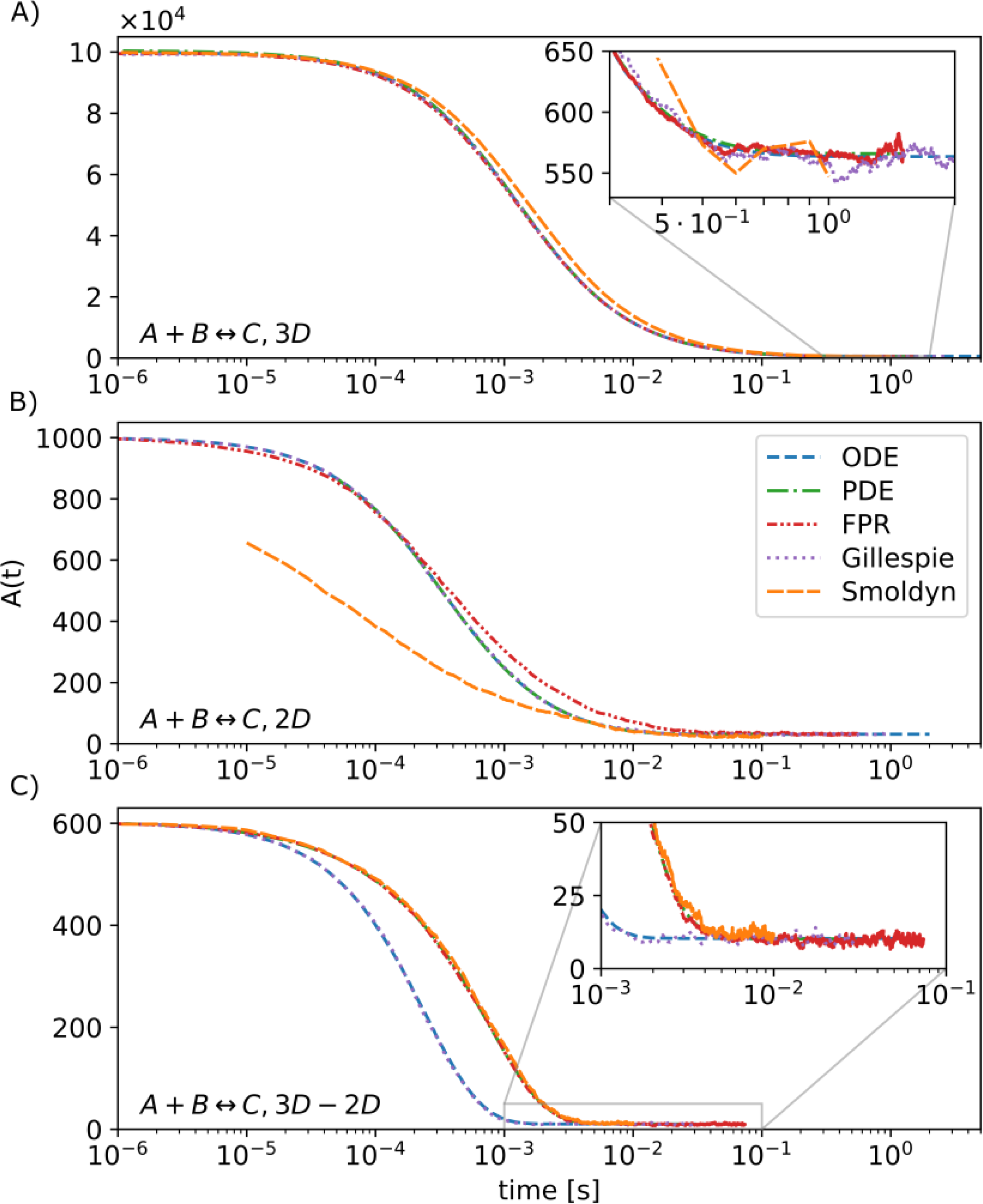
Reversible bimolecular reactions provide fundamental building blocks for any complex biological systems and thus warrant careful testing. In all dimensions, we use well-mixed initial conditions in closed systems, such that the reactions will reach a well-defined thermodynamic equilibrium. We test them using the non-spatial ODE and Gillespie methods, which match each other for all systems. The analytical solution to the rate-equation is essentially exactly captured by the ODE. We compare the spatial PDE solved using VCell with single-particle methods solved via the FPR algorithm (NERDSS) and Smoldyn. For the 3D→2D reaction, we see the first difference between the ODE and PDE solution, because in the spatial models the reactants are well-mixed in distinct locales (volume vs surface), whereas in the non-spatial models, they are all well-mixed in one volume. For 2D, single-particle methods will differ from other approaches, with NERDSS essentially capturing the theoretical diffusion-influenced reaction dynamics in 2D (Yogurtcu and Johnson, 2015).

### 1A: Bimolecular association in 3D, 2D, and from 3D to 2D

Although seemingly simple, bimolecular association events require both a diffusional encounter and a reactive event, thus the rate-constants and the kinetics are dependent on the dimensionality of the systems, and even for well-mixed systems, spatial details can cause deviations from non-spatial models. For reversible bimolecular association of well-mixed reactants in a closed system, the equilibrium is theoretically well-defined and the kinetics for non-spatial rate equations can be derived analytically. One may note that when bimolecular association is reversible, recovering the proper equilibrium is a simple test that can nonetheless be challenging for single-particle methods. Reaction probabilities and the placement of reactants upon un/binding events must be derived to ensure equilibrium is reached (Box 5).

In 3D, all models and tools produce nearly identical results, even for this strongly diffusion-influenced reaction (k_on_=1.48 10^7^ M^-1^s^-1^). This is as expected (Fig. 3a). Although at short times the kinetics is slightly faster for Smoluchowski type simulations (NERDSS), the kinetics rather rapidly converges to the macroscopic rate equations (Johnson and Hummer, 2014). Differences between single-particle and non-spatial methods can also emerge for reversible reactions as they approach equilibrium (Mattis and Glasser, 1998; Tauber et al., 2005), but these can only be effectively observed with high numerical precision and statistics—usually they are dwarfed by the copy number fluctuations.

Unlike in 3D, macroscopic rate equations in 2D only approximate the dynamics captured in Smoluchowski-type approaches at all times (Fange et al., 2010; Hellander et al., 2012; Yogurtcu and Johnson, 2015) (see Box 4). All macroscopic rate-based methods produce the same kinetics as each other (Fig. 3b). Here we see distinctions between the spatial PDE and the spatial single-particle methods. Although species diffuse in the PDE, because they are present at all positions in space (due to uniform initial conditions), association is not dependent on their spatial distribution. For single-particle methods, there is always a distribution of starting separations between species that leads to some very fast reactions initially, and at long times produces slower reactive collisions as particles that started off close to each other have already been consumed in the reaction. Because Smoldyn approximates the dynamics of the Smoluchowski model, the kinetics can be off for specific parameter regimes, with deviations being typically very small in 3D but significant in 2D.

For binding between 3D particles and 2D particles (relevant for biological cases where soluble cytoplasmic proteins bind to membrane proteins or lipids), all models produce the same equilibrium, but the spatial models have slower kinetics delayed by diffusion to the surface (Fig. 3c). The extent of divergence between the non-spatial and spatial models is driven by three factors, the ‘height’ the solution volume stretches from the membrane surface, the diffusion coefficient of the 3D particles, and the speed of the binding. Here we simulated a fast, strongly diffusion-influenced reaction (8.4×10^7^ M^-1^s^-1^), meaning nearly every collision results in a reaction. For a simulation box with *h*=0.2 *µ*m, one can estimate an average time to diffuse to a surface particle would be ∼60 *µ*s (D^3D^=30 *µ*m^2^/s), which is relatively fast. However, with the numerous 2D particles mixed in the solution volume for a non-spatial simulation, the time to diffuse to a ‘surface’ particle drops to ∼6 *µ*s. Thus we find a mean relaxation time of ∼200 *µ*s without space, vs ∼700 *µ*s with space (Fig 3c). By dropping the reaction rate to more moderate protein-protein interaction levels, the spatial and non-spatial results begin to converge. Smoldyn shows excellent agreement for larger steps, here 10^−6^ s, although the kinetics shift slower for shorter steps. We note that when particles can only collide with one another from one side (because one is embedded in a surface, for example), this reduces the binding by a half, and solvers should explicitly account for this so that user-defined rates produce the equilibrium expected from a non-spatial model.

Lastly, transitions to the surface can be modeled using adsorption, which uses an effectively 1D rate. This is more efficient but importantly, it need not reach the same equilibrium as explicit particle simulations, because the occupancy of surface binding sites is not accounted for. Modeling explicit particles thus gives more control over the surface properties, and algorithms for binding to surfaces while accounting for site occupancy using implicit sites rival adsorption models in speed (Fu et al., 2019). Not all tools allow for all types of surface binding, hence it is important to recognize these distinctions between adsorption vs single-site binding.

### 1B: Crowding

Inside of living cells, the extremely high density of macromolecules (with typical spacing on the order of nanometers) can alter the speed of molecular diffusion(Ando and Skolnick, 2010) and kinetics of intermolecular reactions, either increasing or decreasing biochemical reaction rates as compared to rates in dilute solution, depending on the size and mobility of the crowders (Schreiber et al., 2009; Zimmerman and Minton, 1993). As crowders become larger and more immobile (e.g. vesicles) they act more like barriers to encounters, slowing down rates of association (Minton, 2006; Zhou et al., 2008). However, as we see here, when crowders are of comparable sizes and similar mobility to the reactant species, they drive up rates of association. This rate increase is due to a reduction in the total volume available to the reactants, effectively concentrating them without providing substantial barriers to encounters (Minton, 2006; Zhou et al., 2008). To quantify how increasing concentrations of crowding agents alter bimolecular association rates, we here simulated the bimolecular reaction A+B→B+C in the presence of additional, inert, C crowders, where all species diffuse and exclude volume (Fig. 4a). The analytical solution for no crowding/no excluded volume, *A*(*t*) *= A*_O_exp *(-k*_*macro*_ *B*_*tot*_), provides a convenient baseline and fit function for interpreting deviations due to crowding/excluded volume (Fig. 4b).

**FIGURE 4.**
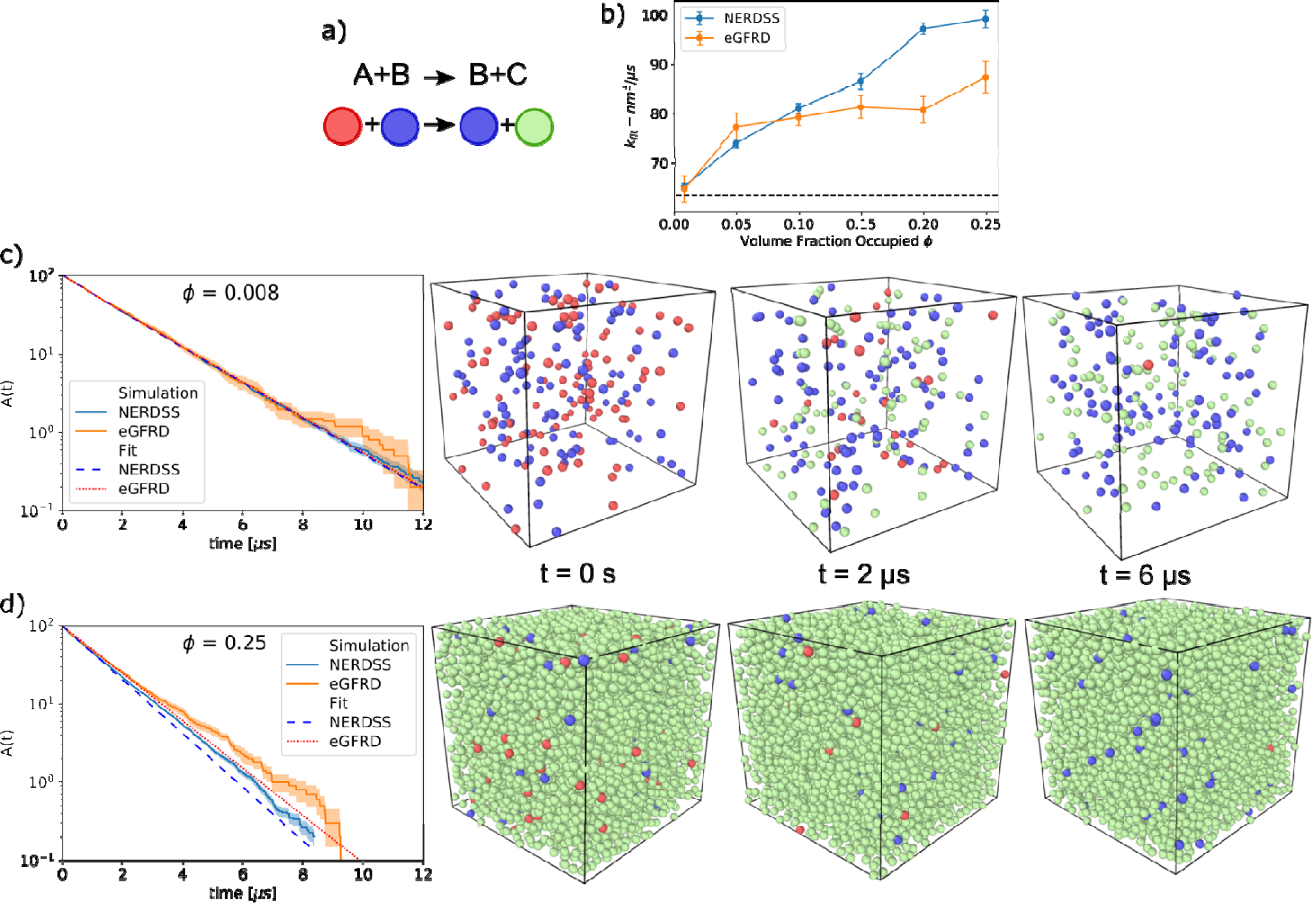
The effects of volume exclusion, or crowding, can be tested by a simple model of bimolecular association in the presence of inert particles. a) For the simple model B+A→B+C, the total population of catalytic B molecules (blue) remains fixed. A molecules (red) are converted into C molecules (green) when they collide with B molecules. C molecules do not react, but do exclude volume and so can act as physical crowders. A molecules (red) are depleted over time. b) The kinetics of A depletion depend strongly on the initial concentration of inert crowders C. For each simulation, the rate of depletion of A is fit to a single-exponential based on the solution to the rate-equation (see Methods). The best fit rate is plotted here as a function of the volume fraction occupied by all A, B and C particles. c) For low crowding fraction where the initial concentration of C molecules C_0_=0, the kinetics is well described by a single exponential with a rate close to the non-spatial solution of 63.5 (65.4 FPR, 64.9 GFRD). d) For high crowding with a large initial C_0_, the simulated kinetics is not as well described by a single exponential, exhibiting slower decay as the density of A approaches zero. Snapshots display the actual volume of the particles (r=0.5nm) given the full volume (23.2nm boxlength), with 100 initial A particles (13.3mM), 100 initial B particles (13.3mM), and variable C. The walls use periodic boundary conditions. Simulations were performed with the NERDSS and eGFRD software, using the FPR and GFRD algorithms, respectively.

For the single-particle algorithms that capture excluded volume (GFRD (van Zon and ten Wolde, 2005) and FPR (Johnson and Hummer, 2014)), two main results emerge. First, the overall kinetics of association increases with increasing crowding fraction, up from *k*_on_=63.5 nm^3^/µs at zero crowding/no excluded volume (3.8×10^7^ M^-1^s^-1^) to ∼85-100 nm^3^/µs with 25% crowding fraction. This result is consistent across both algorithms, indicating that for the well-formulated Smoluchowski model applied to a strongly diffusion-influenced reaction, small mobile crowders will enhance reaction rates for concentrated reactants (here A_0_=13.3mM). This same trend was observed for simulations at lower reactant concentrations but comparable rate constants(Kim and Yethiraj, 2009), but differs qualitatively from a recent study which found that crowding reduced reaction rates at the same reactant concentrations studied here(Andrews, 2020). In the most recent study, however, the crowders were completely immobilized(Andrews, 2020), meaning they did not capture thermal motion of macromolecular crowders, but behaved as a rigid and impermeable barrier. Second, for high crowding regimes, we find that the kinetics is not described by a single rate constant (Fig. 4d), whereas at low crowding the results fit extremely well to the non-spatial analytical solution, with a new rate-constant (Fig. 4c). This is perhaps not surprising; for diffusion-influenced reactions, short-timescale kinetics is dominated by reactants that are already close together, where we expect crowding agents to promote their repeated collisions. Then, as the reactant populations decrease either to zero or towards an equilibrium state, the kinetics slows relative to non-spatial predictions, although this shift may be hard to detect (Johnson and Hummer, 2014; Mattis and Glasser, 1998; Tauber et al., 2005; Yogurtcu and Johnson, 2015). Our results suggest that crowding agents exacerbate this slow search for the final reactants, similar to what happens in 2D, making the deviations from a single-rate constant kinetics easier to detect.

Although mobile crowders do impact reaction rates, the changes are often quite modest. For the strongly diffusion-influenced reaction simulated here, we measure clear increases in rates, but for more rate-limited reactions (*k*_on_=6×10^5^ M^-1^s^-1^), the effect of crowders on the reaction rate is minimal (data not shown). Finally, FPR and GFRD are not in perfect agreement in terms of the quantitative size of the change in kinetics, although qualitatively they both predict a higher rate. At these high densities, GFRD converts to a brute-force Brownian Dynamics algorithm, rather than its exact event-driven method. FPR must also take extremely small time-steps (10^−10^ to 10^−12^ s) to prevent particle overlap. It is not clear which method is truly more correct, as neither will preserve exactly twσbody problems at each step. Further, we note that over the span of a picosecond time-step (10^−12^ s), particle dynamics is not truly diffusive but is inertial. Capturing these dynamics would require a different model (e.g. generalized Langevin Dynamics(Van Kampen, 2007)) that tracked both positions and velocities of particles.

### Category 2: “Intermediate” cases

This category includes slightly more realistic biological cases with interesting emergent properties. These are particularly useful for illustrating the fundamental conceptual differences among the different modeling approaches.

### 2A: Exploiting membrane localization to stabilize protein-protein interactions

In a variety of biological processes, including clathrin-mediated endocytosis and initiation of signal transduction, multi-valent proteins localize to membranes and assemble into larger, multi-protein complexes. Reducing dimensionality from a 3D search to a 2D search can accelerate kinetics of receptor binding (Adam and Delbruck, 1968; Berg and Purcell, 1977), and stabilize interactions for macromolecules restricted to the surface (Abel et al., 2012; Kholodenko et al., 2000; Minton, 1995). For cytoplasmic protein binding partners, localization to the 2D membrane can further drive dramatic increases in protein complex stability, as was recently quantified using a simple model of two proteins that bind to one another and to a specific membrane lipid (Yogurtcu and Johnson, 2018) (Fig 5a,b). The origin of the increased stability is largely a concentration effect, where the reactants collide with one another more frequently on the surface than in solution. While a change also must occur in 3D vs 2D equilibrium constants (Wu et al., 2011), the magnitude is usually much smaller than the change in 3D vs 2D search space (V/A > K^3D^/K^2D^) (Yogurtcu and Johnson, 2018).

**FIGURE 5.**
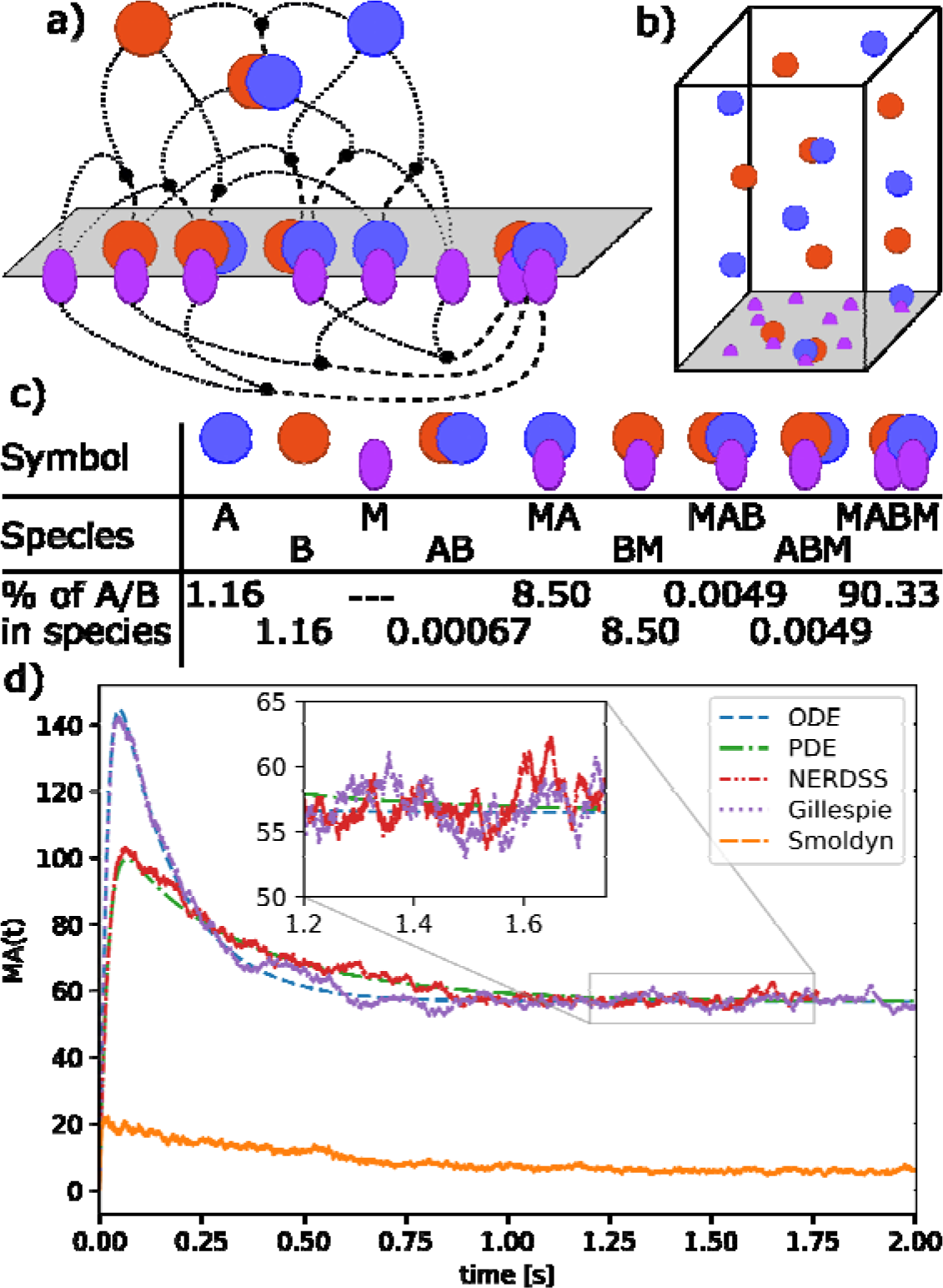
Membrane localization can tune speed and stability of protein complex formation. **a)** A pair of protein binding partners A and B that can also localize to the membrane surface through binding a lipid M can exploit the 2D search space to promote complex formation. There are three binding interactions (below the plane) that involve 2D interactions between protein A and B on the surface, or between a protein and the lipid M. b) The species can partition reversibly between solution and the membrane, forming 9 distinct species, which are listed in (c). c) Model species and fraction of A or B proteins in each at equilibrium. d) Time-dependent formation of a single protein-lipid complex MA. (The time course for BM is identical due to the choice of model parameters.) The initial rise in MA concentration is followed by a drop as the MA and BM complexes combine to form MABM, which dominates at equilibrium. The spatial simulations (PDE and NERDSS) exhibit slower MA formation kinetics and a lower peak concentration than the well-mixed simulations (ODE and SSA) due to slower recruitment of proteins to the surface, which is limited by diffusion. Relaxation of MA to equilibrium is also slower for the same reason. For smaller simulation volumes, where diffusion to the surface is fast, these differences disappear. Smoldyn does not reproduce the proper equilibrium due to inaccuracies of binding in 2D.

With the ability to localize to the membrane and reduce their search space, over 90% of proteins end up bound to one another (Fig. 5c), which contrasts markedly with the purely 3D solution binding result calculated at equilibrium where only 4.6% would end up bound (for K_D_=20 *µ*M and [A]_0_=[B]_0_=1 *µ*M). Although all simulation methods are expected to produce the same equilibrium, and in this example stochastic effects are minimal, we find here that spatial effects arise because diffusion slows the localization of A and B molecules to the membrane (Fig. 5d). The height of the box in the spatial simulations is 5 *µ*m, and so despite efficient solution diffusion (50 *µ*m^2^/s), the proteins are recruited more slowly to the membrane in spatial simulations than in the well-mixed simulations (compare red and green curves in Fig. 5d with purple and blue ones). The protein-lipid complexes plotted in Fig 5d peak before dropping, as they bind to one another to equilibrate, and the peak is lower when they do not localize rapidly to the membrane. Errors in the results calculated using Smoldyn arise due to approximate treatment of purely 2D interactions, which leads to quantitative deviations from expected equilibria and kinetics for these reactions. This current limitation is being actively addressed in Smoldyn software (S. Andrews, personal communication).

Overall, these types of reactions form a critical component of more complex models of membrane-mediated assembly. Due to the quantitative differences we observe here in time-dependence of species numbers, if the model is coupled to reactions that drive it out of an equilibrium steady-state, this could then drive qualitative changes in the biological outcomes (as in case 2B below).

### 2B: Increasing stochastic fluctuations in a system with multiple steady states

Positive feedback in combination with other interactions can give rise to systems with multiple, distinct steady states. Such systems can exhibit large differences in their dynamics depending on whether they are simulated deterministically or stochastically. A simple model that illustrates such effects is the autophosphorylating kinase model first introduced by Lisman (Lisman, 1985) and studied more recently by Agarwal et al. (Agarwal et al., 2012), who analyzed a stochastic version of the model. Figure 6a shows a diagram of the model, which consists of a kinase that can activate itself through phosphorylation (reactions 2 and 3) and a phosphatase that can bind and dephosphorylate the active kinase (reactions 4 and 5). Because the production rate of active kinase (Ap) increases with the amount of Ap starting at low concentrations (Fig. 6b, red curve), the system exhibits positive feedback. The rate balance plot shown in Fig. 6b illustrates a case where the production and degradation rates of Ap as a function of Ap intersect multiple times to give rise to multiple steady states. This model has three steady states, also called “fixed points,” two of which are stable (Fig. 6b, filled circles), For a stable fixed point, the balance of production and decay returns the system back to the fixed point following any slight changes in Ap away from the steady state value, whereas at an unstable fixed point the balance of rates carries the system away from the fixed point as a result of any tiny fluctuation.

**FIGURE 6.**
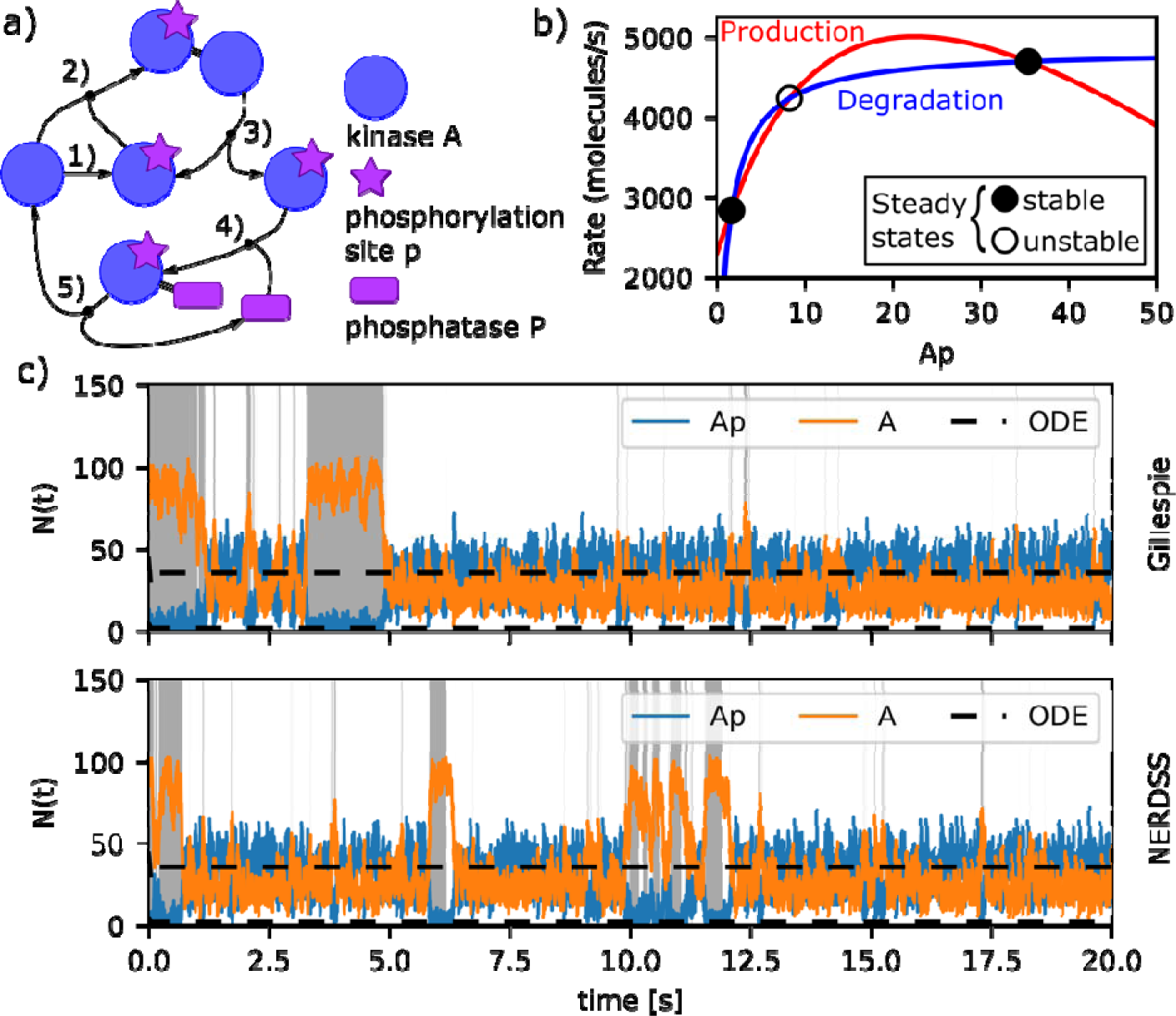
Positive feedback induces stochastic switching in a kinase autophosphorylation circuit. **(a) Model reactions**. The kinase A (blue circle) becomes active upon phosphorylation (purple start; reaction 1) and serves as its own substrate (reactions 2 and 3). A phosphatase P (purple rectangle) binds and dephosphorylates the phosphorylated kinase (reactions 4 and 5). The activation reactions 2 and 3 form a positive feedback for kinase activation. A low rate of spontaneous activation of the kinase (reaction 1) is also included to prevent the system from being trapped in a state with no active kinase. **(b) Rate balance plot identifying steady state concentrations of phosphorylated kinase (Ap)**. The red and blue lines show the rates of production (sum of rates of reactions 1 and 3) and degradation (rate of reaction 4) of Ap for the parameter values and initial concentration simulated here (Methods). Intersections of these curves indicate points at which production and degradation rates are equal and hence give rise to a steady state of the system. The two intersection points shown with filled circles indicate the stable steady states of the system, which occur at Ap concentrations of 1.7 and 35.5 molecules respectively. A third steady state (open circle) indicates an unstable steady state, which occurs at a value of 8.3 Ap molecules. **(c) Model trajectories computed with deterministic and stochastic methods**. Deterministic trajectories starting from different initial conditions may relax to either of the two stable steady states (black dashed lines). Stochastic trajectories exhibit fluctuations about each of the steady states and occasionally switch between states with low and high kinase activation as shown by the blue trajectory. The top panel shows the result of a non-spatial stochastic simulation (Gillespie) and the bottom panel shows the result of a spatial stochastic simulation using the same paramenters (NERDSS). Regions of the trajectories shaded gray indicate where the system is in the state with low kinase activation with correspondingly high levels of inactive kinase (orange lines). In the regions with no shading the system is in a state with high kinase activation, as indicated by the active kinase level (blue lines) generally being above the inactive kinase level. The spatial stochastic simulations performed with NERDSS exhibit small differences from the non-spatial simulations carried out using Gillespie when performed in 3D with all species having a diffusion constant of 100 µm^2^*/*s, and the same macroscopic rate is targeted (see Methods for full details on parameters used in simulation).

Deterministic simulations of the system, whether spatial or not, converge rapidly to one of the stable steady states depending on the initial level of Ap (Fig. 6c, black dashed lines). Simulations starting from a state with low initial Ap will reach the steady state with lower Ap and vice versa for high Ap. Stochastic trajectories, on the other hand, may initially stay in the vicinity of the closer steady state, but fluctuations due to noise occasionally induce switches between states. In both trajectories shown in Fig. 6c the system starts in the lower steady state (as indicated by gray shading) but after a minute or so switches to the higher steady state where it continues to display fluctuations, some of which lead to short-lived excursions back to the lower state. For the set of rate parameters we used here, the system spends more total time in the higher state than the lower state (88% vs. 12%). In addition, the length of each segment in the higher state, called the residence time, is longer on average (2.6s vs 0.34s) (see Methods and Fig S1).

Despite the dramatic differences between the deterministic and stochastic simulations for this model, we find the addition of space has relatively modest effects, with spatial models still producing bistable switching as diffusion slows. Fig. 6c shows a non-spatial SSA simulation in the upper panel (Gillespie) and an explicitly spatial simulation in the lower panel (NERDSS). For the geometries and diffusion constant values chosen here, these two simulations yield small differences between the probabilities and residence times for each state (Table S4). When the diffusion constant is reduced by a factor of 10 the probabilities of the lower state drop significantly although not dramatically, and the residence times of both states drop (Fig. S2 and Table S4). The shorter residence times in spatial simulations relative to the non-spatial likely results from larger fluctuations in copy numbers due to transient spatial inhomogeneities, making both steady-states slightly less stable.

Reduction of the diffusion coefficient by another factor of 10 makes simulation incompatible with the macroscopic reaction rates specified in the model. In other words, the reaction rate is limited by diffusion (see Box 4). We found that the effect of varying diffusion constant was quantitatively similar in the same model, but with all rates reduced by a factor of 10, reducing the fastest rate from 8*10^8^ to 8*10^7^M^-1^s^-1^ (Table S5). A study on a similar model of bistability found that slow diffusion limited the parameter regimes where one observed bistable switching due to fluctuations in local concentration, but also by changing the effective rates(Endres, 2015). Here we kept the macroscopic rates the same as the non-spatial model even as diffusion constants slow, and our results therefore indicate that timescales are sensitive to spatial inhomogenities that persist longer with slower diffusion.

### Category 3: “Application” cases

Proper implementation of the test cases in this category requires the building blocks described above. These well-defined problems yield rich, biologically-interpretable outputs that are sensitive to parameter choices.

### 3A: Stochastic effects in gene expression

Proteins that are important for controlling circadian clocks exhibit regular oscillations in expression levels. These oscillations are quite robust even in the presence of stochastic copy-number fluctuations. In fact, stochastic fluctuations can support oscillations in regimes that a purely deterministic model cannot (Fig. S3). Using a simple model of circadian oscillations (Vilar et al., 2002) (Fig 7a), we simulated the behavior of an activator protein A and repressor protein R that are produced from mRNA transcribed from a single copy of a gene (one for each protein). Coupling of A and R expression is driven by positive feedback of the activator A, which binds to each gene’s promoters to enhance transcription. Protein R also binds to A to effectively degrade it, and all proteins and mRNA are also degraded spontaneously at a constant rate. If the spontaneous degradation rate of protein R is slow, the oscillations will quench in the deterministic model, but persist in the stochastic solutions, as demonstrated in the original work and reproduced in Fig S3.

**FIGURE 7:**
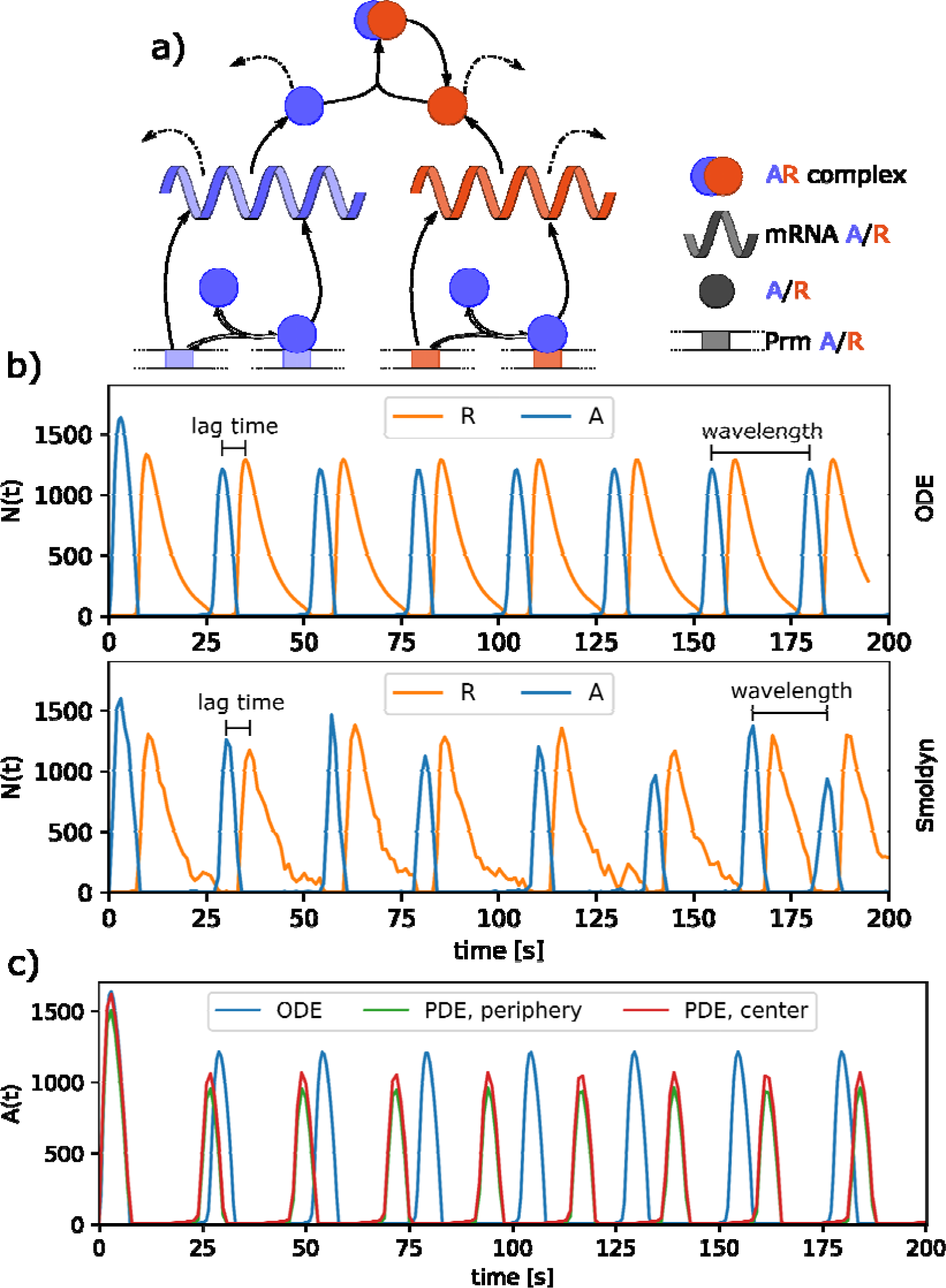
Circadian Clock model shows robustness of oscillations to stochastic fluctuations. a) Model of two proteins A and R, translated from corresponding mRNA and transcribed from the corresponding genes (Prm A/R). Protein A is an activator that binds to the promoter of both genes and increases their transcription. Protein R acts as a repressor that binds to A and catalyzes its degradation. All proteins and mRNA also undergo spontaneous degradation with a specific lifetime (dashed arrows). b) Growth of A copy numbers (dashed) peaks and then decreases as it is depleted by repressor R. Initially most R molecules are present in the AR complex, but the number of unbound R molecules increases as A is degraded. Next, the amoung of free R peaks and then begins to decrease because not enough A is present to promote its transcription at a rate that is greater than its rate of spontaneous degradation. The cycle then restarts, with the ODE solution producing highly regular oscillations of 25.2 s between A peaks, and 6s lag between A and R peaks. The stochastic single-particle simulator Smoldyn produces noisier oscillations, but the average behavior is very comparable to the deterministic ODE model, with peak oscillations at 25.9 s and 6.1 s lag between A and R peaks. c) After localization of the promoters to the cell center and slowing of D_A_ from 10 to 2 *µ*m2/s, the A molecules become more concentrated in the cell center (red vs green), which leads to noticeably faster oscillations relative to the non-spatial solution (blue).

We find that with the addition of space to the model, with all species diffusing at *D*=10 *µ*m^2^/s, the oscillation times show no real significant differences in single-particle or deterministic solutions relative to the non-spatial model (Table S6). To quantitatively compare the kinetics across all models, we cannot use simple steady-state values, due to the oscillations. These time-dependent oscillations are nearly perfectly regular in the deterministic models, but are quite imperfect (although recurring) in all stochastic and single-particle methods (Fig. 7b). We therefore measure the average time interval between peaks in the expression of A, and the lag time between the appearance of a peak in A expression followed by a peak in R (see Methods). The similar results across all methods shows how with relatively small spatial dimensions (sphere of R=1 *µ*m), purely 3D reactions, and all species diffusing at the same *D*=10 *µ*m^2^/s, no spatial dependence was distinguishable.

The lack of any significant spatial dependence is somewhat surprising, because we pushed the reaction rates into the strongly diffusion-influenced regime. The model was actually formulated to describe the slow oscillations of gene expression regulation, with all rates reported in units of hr^-1^, rendering diffusion times irrelevant. Here, we chose to accelerate the rates by a factor of 3600 (from hr^-1^ to s^-1^), due to the computational expense of simulating spatial models with explicit diffusion. This result overall indicates that the reactants still mixed faster than any spatial inhomogeneities could emerge, preventing deviations between spatial and non-spatial simulations.

We were finally able to measure a significant dependence on diffusion when we simulated the same set of reactions but we localized and immobilized the promoters for A and R to a small nucleus in the center of the volume, with unrestricted diffusion in and out of this nucleus (Fig 7c). For the same diffusion constants of *D*=10 m^2^/s for all other species, this immobilization of promoters was not enough to have an effect, and oscillation times remained not significantly different (Fig S4 and Table S7). However, when the diffusion constant of the activator A was slowed to 2 m^2^/s, we observed a persistently higher concentration of A and its mRNA near the cell center relative to the cell periphery (Fig 7 and Fig S4). This localization of A near the gene promoters had the effect of shortening the oscillation period in the PDE from 25s to 22.5s (Table S7). This change was independent of the diffusion constant of R, which does not bind the genes. Further, this same trend was observed in the single-particle simulations, which also produced faster oscillations as D_A_ slowed to 2 m^2^/s (Table S7). Overall, we thus found that while slowing the search time for the Activator for both genes does modulate the oscillation times, the oscillations are quite robust and strongly dictated by total copy numbers and the unimolecular decay reactions, which are inherently independent of diffusion and spatial dimensions. This fairly complicated scenario is one where intuitively we had expected to see strong spatial effects, so it is rather interesting that the actual spatial effects are minor in this case. Certainly this outcome might vary depending on the parameter regimes selected.

### 3B: Spatial and temporal oscillations in MinCDE

When coupled to reactions, diffusion that is sufficiently slow or occurs over large enough length scales can establish spatial gradients of concentrations and subsequent oscillations in space and time. By construction, spatial oscillations are lost in non-spatial models, and no temporal oscillations occur either (Fig S5). In a simplified model of bacterial cell division, the MinD and MinE proteins (Huang et al, PNAS 2003) spatially control the location of cell division by creating an oscillating spatial gradient of both proteins in the cytosol and on the membrane (Fig 8). The site of cell division is determined by assembly of a ring constructed from the bacterial tubulin homolog FtsZ. Because the Min proteins inhibit the assembly of the FtsZ ring, the spatial oscillation of the Min proteins from one end to the other in the cylindrical bacterial cell results in the localization of the FtsZ ring to a site very close to the geometrical center (Raskin and de Boer, 1999). This model involves 3D and 3D→2D bimolecular reactions, and unimolecular reactions, with no 2D reactions. Instead of forming explicit polymers, the membrane-bound proteins act to locally increase protein density through recruitment.

**FIGURE 8.**
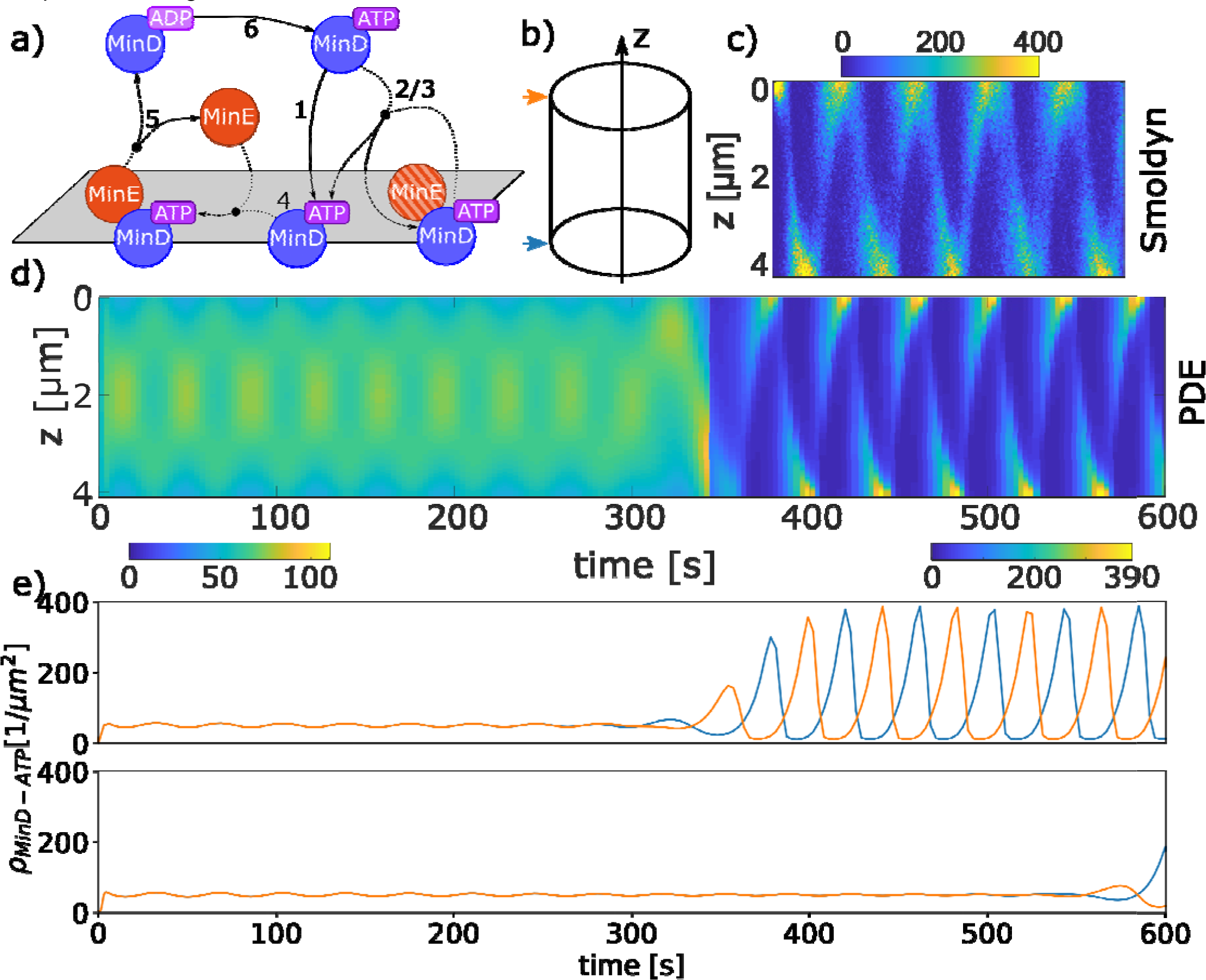
Spatial oscillations in protein abundance for the MinCDE cell division model. a) The MinD protein can exist in two states, ADP or ATP bound (named MinD-ADP and MinD-ATP). Only MinD-ATP localizes to the membrane in a 3D→2D reaction (producing MinD-ATP^2D^). Once on the membrane MinD-ATP^2D^ can recruit additional cytoplasmic MinD-ATP (3D→2D) or MinE (3D→2D). The MinD-ATP.MinE^2D^ complex on the membrane recruits MinD-ATP from solution as well (3D→2D), or it dissociates to return MinE to solution and MinD-ATP^2D^ to MinD-ADP. None of these 6 reactions is thus reversible. b) Simulations in a cylindrical cell with L=4um and R=0.5um, with 2.1143*µ*M MinD-ATP and 0.74*µ*M MinE initially well mixed. c-d) The kymographs show how the copy numbers of minDt on the membrane in molecules/µm^2^ oscillate in space and time. c) single-particle simulation Smoldyn d) PDE with uniform initial concentrations has faint symmetric oscillations visible up to ∼300s before symmetry breaks. e) Time-dependence of MinD-ATP^2D^ at a point on the left (blue) and right (orange) end of the cell. The oscillations are perfectly symmetric until ∼300s in the upper panel. However, if the error tolerance is tightened on the numerical integration, the symmetric oscillations persist longer out to ∼550s in the lower panel, illustrating the dependence on numerical precision.

We are able to produce very similar spatial and temporal oscillations of the MinDT protein on the membrane in both a stochastic single-particle model (Smoldyn (Andrews, 2017)) and a deterministic PDE solution (using Vcell (Moraru et al., 2008; Schaff et al., 2016)). The major distinction between the two models is that the deterministic PDE is able to support symmetric, or striped oscillations in the protein, as would be generated by two traveling waves in opposite directions. However, we find that these dynamics are not stable even in the deterministic solution—the accumulation of numerical precision error eventually breaks the symmetry and transitions to pole-tσpole oscillations. We show that by increasing the error tolerance on the numerical integration or by increasing the PDE mesh size one can delay the onset of this transition, clearly demonstrating the dependence on numerical precision (Fig 8e and Fig S6). The stochastic single-particle method is therefore not able to support the symmetric oscillations due to copy number fluctuations, and always produces pole-tσpole oscillations (Fig 8b). This shows quite clearly that the pole-tσpole oscillations are more robust to stochastic fluctuations in copy numbers for this geometry, and in the wild-type biological systems, these are the types of oscillations always observed for cells of this length (Raskin and de Boer, 1999).

A challenging aspect of model comparison for any simulations tracking the spatial and time-dependent concentration of species is deciding how best to quantify the results. Analysis is often defined in a problem-specific way. For the MinCDE model, depending on the cell geometry and model parameters, it may not even reach a steady-state over long time scales, but can continue to oscillate and change oscillation amplitude or frequency. For symmetric geometries (like cylinders), we reduced the analysis to a function of time by tracking the MinD-ATP^2D^ molecules at a specific point on the membrane (Fig S7). Then, similar to the analysis for Fig 6, we calculated the average period between oscillations (Methods), finding values of 43.0±0.7s for Smoldyn and 41.3±0.3 s for the PDE.

The oscillations in this model are fairly sensitive to initial concentrations of species (Fig S6) and their relative stoichiometry. If the initial concentrations were cut in half, we found the oscillations disappeared in both models. The MinCDE model has been extensively studied, with the major determining features of oscillations being initial concentrations and cell geometry (see (Halatek and Frey, 2012) and ref therein). The oscillations are relatively robust even to small changes in the reaction network, as long as membrane recruitment (3D→2D) and ATP hydrolysis are included (Halatek and Frey, 2012). We note that for Smoldyn, the oscillations were sensitive to the diffusion coefficients of the membrane bound particles, set here to D=0.05 *µ*m^2^/s. If proteins did not diffuse on the membrane, the oscillations disappeared in Smoldyn, whereas they were unaffected in the PDE. This is in part because the molecules do not have excluded volume in Smoldyn, and because no explicit polymers were formed, they accumulated in a small space rather than spreading along the membrane.

### Summary of Test Case Outcomes

By simulating the same biological cases using a range of spatial, non-spatial, stochastic, and deterministic methods, we have shown here how specific biological features and in some cases algorithm selection (e.g. classes of single-particle approaches) can alter quantitative and and even qualitative outcomes. As summarized in Fig 9, quantitative differences can be small, medium, or large based on system geometry and parameter regimes. Qualitative differences can be minor, but can also be major, producing entirely new behavior patterns that emerge. For example, major impacts could be observed in application scenario A2, which combines an elongated geometry with a comparatively slow diffusion (in comparison to the reaction rate), and includes both 3D and 2D dynamics. In general we find that effects of introducing stochasticity to a system may be orthogonal to its spatial properties, as wrongly applying a deterministic method is likely to dominate all other choices. This can be most clearly seen for the results as shown in I2. Overall, we found that, while stochasticity played a minor role in models with reversible interactions (U1, I1), it was capable of driving major changes in models with irreversible interactions and feedback (I2, A2). Quantitatively, crowding might have a large impact, in particular if combined with slow diffusion and high reaction rates. Also the impact on dynamics that involve reactions in 2D, be this 3D→2D or 2D dynamics, is significant and cannot be ignored in many cases. The suitability of specific simulation methods and their limitations have also to be taken into account when any investigator is making choices about which methods to employ for a particular biological problem (see Table S2). For example, if crowding needs to be considered, particle-based approaches, in particular those of the Smoluchowski type, should be applied. However, in the case of very dense crowding, these approaches do not scale because of their computational cost.

**FIGURE 9.**
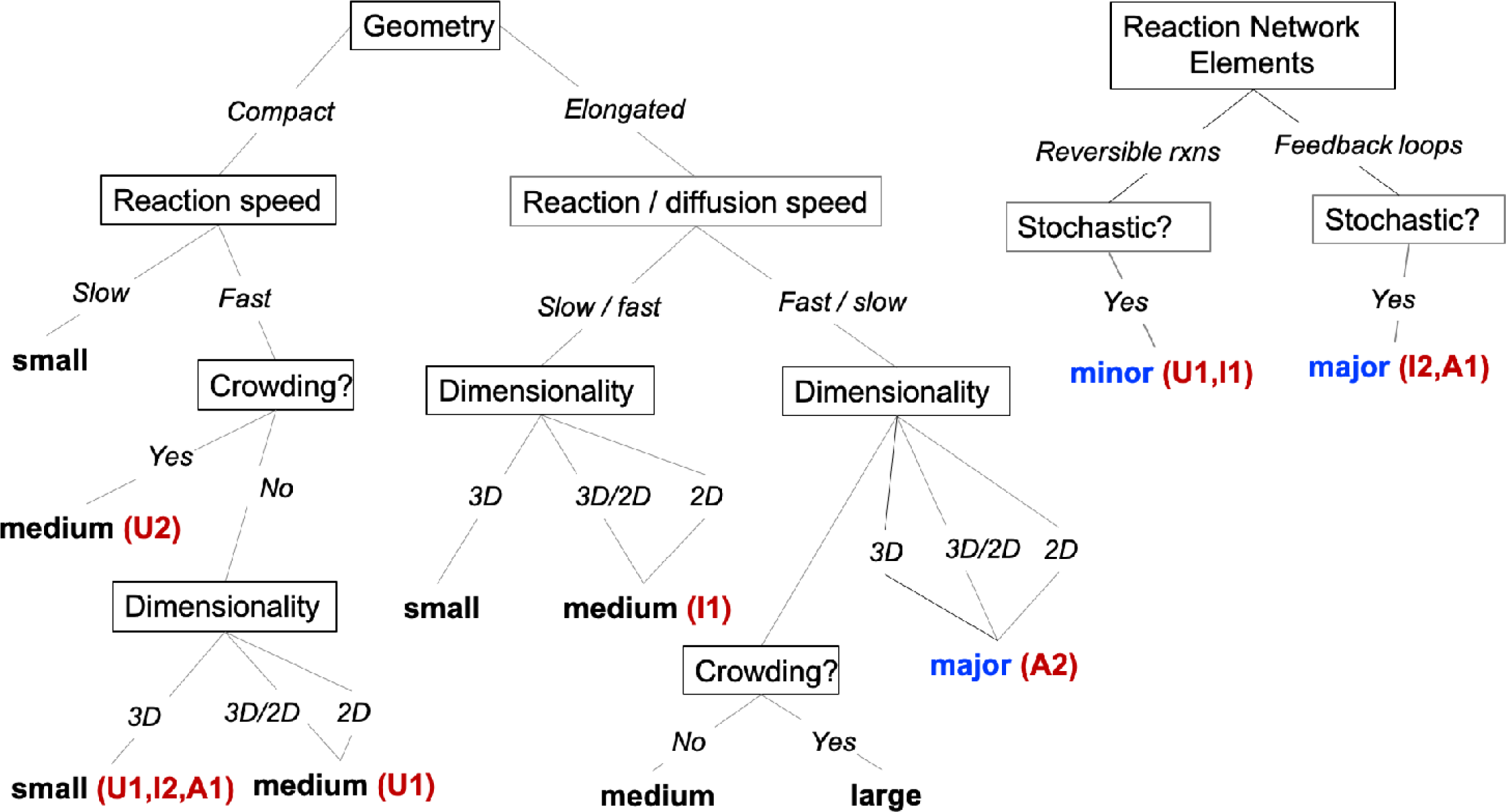
Summary of impact of spatial modeling and simulation approaches on quantitative (small, medium, large) and qualitative (major/minor) biochemical behavior. For quantitative behavior, our tree structure separates parameter and geometry regimes to specifically identify the scale of change observed and expected. For qualitative behavior, we note that, especially for reaction-network elements and stochastic effects, observing major changes also depends on additional parameter specifications, but in a less predictable way than the spatial effects (e.g. relative sizes of distinct rates or copy numbers). Minor changes is the default, as we observe for models with purely reversible reactions (rxns). The test cases range from very simple problems (U1 *Bimolecular association in 3D, 2D, and from 3D to 2D*, U2 *Crowding*), via intermediate tests (I1 *Exploiting membrane localization to stabilize protein-protein interactions*, I2 *Increasing stochastic fluctuations in a system with multiple steady states*) to applications that combine different spatial features (A1 *Stochastic effects in gene expression*, A2 *Spatial and temporal oscillations in MinCD*E). Various tools have been applied in the test cases: U1--ODE, PDE, SSA/Gillespie, particle-based (NERDSS, Smoldyn); U2 – particle-based (NERDSS, eGFRD); I1 – ODE, PDE, SSA /Gillespie, particle-based (NERDSS, Smoldyn); I2 -- ODE, SSA /Gillespie, particle-based (NERDSS); A1 – ODE, PDE, SSA /Gillespie, particle-based (NERDSS, MCell, Smoldyn); A2 – PDE, particle-based (Smoldyn). A detailed description of the theoretical basis and the features offered by these tools/methods can be found in Table S2.

The spatial dimension in our test cases primarily drives quantitative changes in outcomes of varying magnitude. However, once multiple features are combined, as in the A2 MinCDE model, major qualitative differences emerge. We expect that our results will extrapolate to informing model selection in a broader range of biological processes. For example, the effect of clustered molecules acts to spatially localize interactions, which similar to the membrane localization (I1) can drive dramatic increases in complex stability, and changes to kinetics. Compartmentalized or highly elongated and narrow geometries also can act to locally alter species concentrations, not unlike the role of dimensionality in driving quantitative changes (U1, I1) or qualitative changes (A2). Lastly, in all the models studied here, all species were initially well-mixed in their system volumes, albeit in U1 (3D→2D) and I1 there was both a membrane and a solution volume. By introducing an explicit spatial concentration gradient via localized sources or sinks of molecules, spatial effects are inevitable, and can only be captured with explicit spatial representations.

## DISCUSSION

Since the use of GFP as a genetically encoded fluorescent tag came into widespread use over a quarter-century ago (Chalfie et al., 1994), methods for live-cell light microscopy have advanced spectacularly, and the spatial resolution and temporal resolution now possible for imaging experiments in living cells is truly awe-inspiring (Chen et al., 2014; Guo et al., 2018; Liu et al., 2018). At the same time, a great deal of technical development has gone into the generation of live-cell fluorescent reporters for biological readouts including signaling events, enzymatic activity, and force generation (Greenwald et al., 2018; Yasunaga et al., 2019). Many of the cell biological phenomena illuminated by direct observation of dynamic molecular events in living cells reveal a spatial component that could not be fully appreciated during the era when the most sensitive assays for molecular events were biochemical in nature (Machacek et al., 2009; Tay et al., 2010).

At this point, the quality of time-resolved live cell imaging data for cell biological events, and its quantitative accuracy, is far ahead of our ability to model these events computationally. As a community we could avoid doing the work to develop detailed, spatially resolved models when the data quality was relatively poor, relying on images of fixed cells at poor spatial resolution, but now that the data is excellent there is no excuse. Quantitative data demands quantitative models.

Accurate, spatially-resolved 3D modeling of subcellular processes is inherently challenging, particularly as compared to the larger field of time-resolved cellular modeling where space is not taken into account. In exploring the currently available tools for 3D modeling, and dreaming about what the future might hold, we have identified several sticking points that will require focused community effort to overcome.

### Spatially accurate modeling is intrinsically computationally expensive

Most interesting cell biological events unfold over minutes, hours or days, requiring that relevant computational models be able to simulate comparable lengths of real time. For spatially resolved models, keeping track of the positions as well as the numbers and identities of relevant molecular species requires substantially greater computational overhead than simpler models that include only numbers (or concentrations) changing over time. In practice, this difficulty often limits spatially-resolved models to keeping track of only a few molecular species.

An interesting analogy can be drawn to the field of molecular dynamics simulations, which aims to model relative movements of atoms within proteins (Dror et al., 2012), while models of the scales that we are discussing typically aim to simulate the relative movements and transformations of molecules within cells. The equations of motion for atoms within protein molecules are relatively well understood, but accurately calculating their trajectories over even the millisecond time scales relevant for protein folding or conformational changes requires massive computational power (Shaw et al., 2010). Generally, increases in complexity and time scale for simulations of atoms in proteins have benefited from two types of efforts, improvements in hardware including purpose-built computers and distributed computing architectures for parallelization (Snow et al., 2002), accompanied by improvements in statistical sampling approaches to capture rare but significant events (Zuckerman, 2011). Even so, most molecular dynamics simulations still cannot simulate processes central to the lives of proteins such as enzymatic catalysis, because they essentially treat atoms as hard balls subject to defined forces, but ignore the quantum mechanics that would be required to describe chemical transformations. Some hybrid methods have shown promise by including quantum-mechanical detail only in the enzyme’s active site (Karelina and Kulik, 2017), but naturally this improvement in chemical detail comes at substantial computational cost.

Similarly, for spatially resolved models at the cell biological scale, there is room for substantial improvement in efficient design of hardware and computational parallelization, development of appropriate statistical sampling methods, and hybrid methods that could smoothly integrate full detail in some particular locations or at some particular times with computationally efficient approximations elsewhere (Hellander et al., 2017). GPU architectures offer many advantages for efficient parallelization, and have been effectively used by several groups to push cell biological models to much larger scales (Ghaemi et al., 2020; Hallock et al., 2014), though these approaches have not yet been widely adopted. There are also some kinds of modeling problems that intrinsically cannot be parallelized, where each step in time depends on the state of the system as a whole (Enculescu et al., 2010).

Several features of cell-scale modeling render this a massively more difficult computational challenge as compared to molecular dynamics. One, of course, is the scale; a protein may include thousands or tens of thousands of atoms, while a typical cell will include 2-4 million proteins per cubic micrometer of volume (Milo, 2013), or over 10^10^ in an average mammalian cell, not to mention lipids, nucleic acids, carbohydrates, small metabolites and ions that may all be relevant to propagation of cellular signals. At present, all spatially-resolved models for cell biology therefore necessarily include only a very small sampling of the molecular species that may influence the process. Furthermore, most parameters as fundamental as molecular concentrations and reaction rate constants are not known, or known only approximately (Gutenkunst et al., 2007; Schillings et al., 2015). For that reason, cell biological models must typically also perform a broad sampling of “parameter space”, limiting the depth to which any particular instantiation of a spatially-resolved model can be explored.

### There is no common framework for sharing and reproducing spatially resolved models

Over the past decade, there has been a humbling realization that a large fraction of published results in experimental biology, including many that have led directly to significant efforts toward drug development, cannot be reproduced (Begley and Ellis, 2012; Prinz et al., 2011). This has led to widespread reform in both the performance and the publication of experimental research, notably with the US National Institutes of Health (NIH) now requiring explicit training in experimental rigor and reproducibility, and altering the structure of research grant applications to address these issues directly (Health, 2018). In order for the modeling enterprise to advance, it will be necessary for our community to establish standards for sharing and reproducing spatially resolved computational models as well.

Sharing and reproducing models is only possible if every quantitative parameter is strictly defined in a meaningful way. This will require a common “language” that is flexible enough to describe a wide variety of complex cell biological problems, and it is particularly challenging to define parameters that can be implemented in an equivalent way for both PDE-based and particle-based simulations of the same process. Even within the confines of our small working group, where we had all agreed to work together closely and were in constant communication throughout the process, we frequently found it challenging to run the “same” model on different platforms because of inconsistent parameter definitions. We also found that published models often lack critical parameters even if they appear at first glance to be completely described.

For time-resolved cell biological models that do not include an explicit spatial component, the Systems Biology Markup Language (SBML) represents a large-scale community effort to improve standardization of model representation. An extension to SBML explicitly designed for spatially-resolved modeling, SBML-spatial, is nearing completion (SBML.org, 2020) and is supported to varying degrees by some modeling systems, although the specification has not yet been formally released and is not widely adopted. Some leaders in the field have begun to organize resources to aid in efforts toward verifying simulation results (reproduciblebiomodels.org, 2020). Several related fields that require highly complex computational models, such as finite element modeling in biomechanics, have also begun to define community standards for reporting and verifying model parameters and results (Erdemir et al., 2012).

Minimal steps toward ensuring reproducibility in cell biological spatial modeling must include a general expectation that all code will be readily accessible (via GitHub or equivalent) and directly runnable by outside users, not dependent on local files or libraries. All code for spatial modeling should include explicitly defined unit tests, such as those we have included here. Furthermore, all parameters used in any model must include meaningful units. This is not a trivial consideration; for example, if the problem being studied involves a transition between a soluble form and a membrane-associated form of a particular molecular species, we have found that rate constants and their units are usually not carefully documented in most published models, and in some cases it is not even clear that that different researchers agree on what the “right” units are for some typical problems.

### It is difficult to compare outcomes of different types of models, and to compare model results to experimental data

A typical goal for cell biological modeling is to determine whether a particular underlying mechanistic hypothesis is consistent with experimental observation. For processes with a spatially-dependent stochastic component, which includes many cell biological problems of interest, each computational run (and, indeed, each independent experiment) will give a slightly different result. In order to compare experimental results with simulations, and to compare outcomes of different simulations with one another, the simplest approach is to use some kind of summary statistics that described probabilities of various outcomes. However, it is not often trivial to determine which summary statistics are most appropriate. For example, one influential early model illustrating the importance of stochasticity in genetic regulation leading to cell fate determination was an exploration of the events underlying the switch for bacteriophage lambda from a lysogenic to a lytic state, published in 1998 (Arkin et al., 1998). However, more than a decade passed before a useful method for calculating summary statistics on these extremely rare switching events was developed (Morelli et al., 2009). Ironically, it may in many cases be easier to repeat an experiment for thousands or tens of thousands of cells, for example using automated videomicroscopy (Cai et al., 2018), than to run a computationally intensive spatial simulation the same number of times. In general, the field would benefit from greater attention to scalability in model construction.

Some statistical strategies that were originally developed in other fields have been creatively adapted for applications in spatially-defined stochastic simulations. These include extensions to the singular spectrum analysis methods that are widely used for analysis of time-series data (Shlemov et al., 2015) and the weighted ensemble sampling method that is useful when rare events are of particular interest (Donovan et al., 2016).

Visualization of model outcomes should also be standardized. This is much more conceptually challenging for spatially resolved models than for models where the only output is a change in concentration of molecular species over time. Here recent developments in the field of visual analytics, which combines statistical analysis with visualization and user interactions to cope with large complex data sets, may offer some promise (Matković et al., 2018). Visual analytics takes the human into the loop for exploring data and the space of simulation experiments. Other approaches aim at testing requirements and confirming behavioral expectations automatically. One example of a promising avenue for automatic analysis would be spatiσ temporal model checking approaches, which rely on explicitly stating spatiσtemporal properties to be checked on the produced trajectories of the spatial simulation (Bartocci et al., 2015).

### It may not be possible to address distinct model features required for different kinds of cell biological problems within a single modeling framework

In our comparative work, we emphasized simple biochemical interactions because we were able to implement these examples in comparable ways across multiple different computational platforms. We chose to use only very simple geometries for our example cases, but some of the stochastic simulation platforms we considered are capable of incorporating complex cell geometries, and complex geometries are also standard in most PDE solvers. However, none of the packages we used here are designed to incorporate correct subcellular physics for cytoskeletal mechanics, force generation, fluid flow, or electrostatics. Clearly these physical elements are important in accurate simulations of cellular behaviors. It has been widely recognized for over a century that cellular organization is tightly coupled to cellular structure (Abbot, 1916).

In our opinion, the highest priorities for integration of correct subcellular physics with stochastic molecule-based methods of the kind described here are accurate mechanical models for cytoskeletal filaments, molecular motors, and membranes. Interactions of the cytoskeleton and its associated motors with membranes determine overall cell shape and organelle localization, and these also underlie interactions of cells with one another and with their extracellular matrix. Actions of cytoskeletal filaments and motors directly deform membranes to produce invaginations or protrusions, and membrane surfaces influence cytoskeletal filament growth both because of their mechanical resistance (Keren et al., 2008) and because of their ability to accumulate and spatially organize key regulators (Mullins et al., 2018). Simulation frameworks that could properly deal with these kinds of mechanical and biochemical interactions should be adaptable for a wide variety of related problems, for example exploring the assembly of viral “replication factories” on intracellular membranes of host cells, a process which requires large-scale oligomerization of viral polymerase (Spagnolo et al., 2010).

While we recognize that all models are necessarily approximations, we believe that some approximations are better than others. For membrane mechanics, the Helfrich model (Helfrich, 1973) is the most widely used framework for calculation of energies associated with membrane bending and deformation (Guckenberger and Gekle, 2017). While the theory itself is simple and elegant, calculation of energies for highly complex geometries in a continuum mechanics model can become complicated. A few interesting approaches have been used to enable cellular geometries to adapt in response to biochemical reactions (Angermann et al., 2012; Tanaka et al., 2015), but so far none of them have attempted to incorporate correct Helfrich-derived bending energies. PDE-based models for membrane-protein interactions can better approximate the correct underlying physics, but have so far been limited to fairly simple geometries (Rangamani et al., 2014; Wu et al., 2018). Ultimately, the membrane mechanics must be coupled to the proteins and protein assemblies that provide most of the force necessary to induce membrane bending and remodeling.

Capturing the dynamics of protein assemblies, both as highly ordered filaments or lattices (e.g. microtubules or clathrin-coated vesicles) and as disordered condensates, will be essential for bridging biochemical interactions with mechanics. Recent advances in extending single-particle methods to include rigid (Varga et al., 2020) or polymer-like structures (Hoffmann et al., 2019; Michalski and Loew, 2016) have enabled simulations of highly ordered clathrin lattices and viral shells (Varga et al., 2020), or disordered condensate-like assemblies (Chattaraj et al., 2019), respectively. For cytoskeletal mechanics, in many cases a simple approximation of cytoskeletal filaments as elastic rods and motor proteins as discrete localized generators of quantized directional force can give impressively realistic simulation outcomes (Nedelec and Foethke, 2007b). This approach has been widely used for studies of dynamic microtubule and actin structures in eukaryotic cells (Karsenti et al., 2006; Odell and Foe, 2008; Rubinstein et al., 2009). It has recently been demonstrated that mechanically accurate modeling of cytoskeletal filament dynamics can be fruitfully combined with a continuum model for membrane bending in order to simulate endocytosis (Akamatsu et al., 2020).

More generally, it is clear that much more work needs to be done in the development of multi-scale simulations and hybrid computational approaches (Groen et al., 2019). For example, in some intermediate regimes approximate subvolume-based simulations may be able to bridge the gap between well-mixed and particle-based simulations. It will require careful work to understand what approximations are generally useful when jumping across scales in hybrid models, and what approximations give rise to propagating errors.

Overall the members of our working group have thoroughly enjoyed this opportunity to critically evaluate the state of tools that are currently available for simulation of cell biological processes. Although we have of course identified important gaps, we are generally optimistic about recent developments and look forward to incorporation of more sophisticated computational methods to improve the efficiency and accuracy of cell biological modeling as these methods are developed. Spatially accurate cellular simulation is a field still in its infancy, but its future is bright.

## METHODS

### Simulation Solvers

ODE, PDE, and Gillespie simulations were all run using Virtual Cell, version 7(Moraru et al., 2008). Unless otherwise noted, the PDE solver used a fully-implicit finite volume, regular grid (variable time step), max step of 0.1 s. Absolute error tol: 10^−9^, relative error: 10^−7^. The mesh used the default values, which typically resulted in mesh side lengths of ∼0.02 *µ*m. We note that for the PDE, the mesh, volume, or concentrations sometimes had to be adjusted slightly, to ensure the copy numbers were the same for accurate comparisons. The ODE used the Combined Stiff Solver (IDA/CVODE), abs error tol: 10^−9^, relative error tol: 10^−9^. Gillespie simulations used the Gibson-Bruck method, an exact stochastic simulation algorithm, implemented in Virtual cell, or for the autσ phosphorylation model, RuleBender(Smith et al., 2012).

All single-particle solvers propagated Brownian dynamics, with no additional forces added. Smoldyn simulations were run either through Virtual Cell, version 7, or using the stand-alone Smoldyn software, version 2.60/2.61(Andrews, 2017). FPR or NERDSS simulations were run using the NERDSS software, version 1(Varga et al., 2020), which uses the FPR algorithms(Johnson, 2018; Johnson and Hummer, 2014; Yogurtcu and Johnson, 2015) to solve the single-particle reaction-diffusion model. GFRD simulations were run using eGFRD software(Sokolowski et al., 2019) from August 3^rd^ 2019. MCell simulations were run using MCell Version 3.3(Kerr et al., 2008).

### Models and simulation parameters

Unless otherwise noted, species in spatial simulations were initialized as uniformly distributed.

#### Units Tests

##### 1a. 3D Reversible binding A+B ⇌ C

Model parameters: V=3.1934 *µ*m^3^ cube, [A]_0_=[B]_0_=52 *µ*M, or 100,000 copies, [C]_0_=0. k_on_=1.476 × 10^7^ M^-1^s^-1^, k_off_=0.02451 s^-1^. Microscopic rates, k_a_=1000 nm^3^*µ*s^-1^, k_b_=1s^-1^. K_D_=0.00166 *µ*M. For FPR/NERDSS, *σ*=1 nm. D_A_=D_B_=D_C_=1 *µ*m^2^/s. [A]_eq_= 0.293 *µ*M, or 563.5 copies, reached after ∼1 s.

##### Sim parameters

Single-particle methods Δt=10^−7^-10^−6^ s. FPR/NERDSS N_traj_=18, Gillespie N_traj_=10, and Smoldyn N_traj_=1.

##### 1b. 2D Reversible binding A+B ⇌ C

Model parameters: A=1 *µ*m^2^, flat surface. [A]_0_=[B]_0_=1000 *µ*m^-2^, or 1000 copies, [C]_0_=0. k_on_=3.07 *µ*m^2^s^-1^, k_off_=3.07 s^-1^. Microscopic rates, k_a_=10 *µ*m^2^s^-1^, k_b_=10 s^-1^. K_D_=0.00166 *µ*M. For FPR/NERDSS, *σ*=1 nm. D_A_=D_B_ =1 *µ*m^2^/s, D_C_=0.5 *µ*m^2^/s. [A]_eq_= 31.1267 *µ*m^-2^, reached after ∼0.1 s. *Sim parameters:* Single-particle methods Δt =10^−7^s. FPR/NERDSS N_traj_=5, Gillespie N_traj_=5, Smoldyn N_traj_=2.

##### 1c. 3D→2D Reversible binding A+B ⇌ C

Model parameters: V=1 *µ*m^3^ cube, flat membrane surface of 2.2 *µ*m x 2.2 *µ*m. [A]_0_=1 *µ*M, or 602 copies, [B]_0_=6045 *µ*m^-2^, or 29,258 copies, [C]_0_=0. k_on_=8.44 × 10^7^ M^-1^s^-1^, k_off_=70.085 s^-1^. Microscopic rates, k_a_^3D^=500 nm^3^*µ*s^-1^, k_b_=250 s^-1^. K_D_=0.83 *µ*M. *σ*=1 nm. D_A_ =30 *µ*m^2^/s, D_B_=1 *µ*m^2^/s, D_C_=0.97 *µ*m^2^/s. [A]_eq_= 10.3 copies, or 0.017 *µ*M, reached after ∼0.02 s. *Sim parameters:* Single-particle methods Δt=10^−7^s. FPR/NERDSS N_traj_=10, Gillespie N_traj_=5, and Smoldyn N_traj_=3. Smoldyn Δt=10^−6^ s, with the longer steps converging to the PDE solution.

##### 1d. Crowding A+B→B+C

Model Parameters: V=12,500 nm^3^ Cube (side length of 23.21nm), periodic boundary conditions enforced. [B]=100 copies (13.3 mM). [A]_0_=100 copies. [C]_0_ varied across 6 systems: [0, 994, 2187, 3381, 4575, 5768], corresponding to total volume fractions occupied of [0.00838, 0.05, 0.1, 0.15, 0.2, 0.25]. Reactions: 1. A+B→B+C k_a_=85 nm^3^/*µ*s, 2. A+A k_a_=0, 3. A+C k_a_=0, 4. B+B k_a_=0, 5. B+C k_a_=0, 6. C+C k_a_=0. With *no volume exclusion*, Rxn 1 has k_on_=63.5 nm^3^/*µ*s = 3.82 × 10^7^ M^-1^s^-1^. *σ*=1 nm, and thus each particle is effectively modeled as a volume-excluding sphere of diameter=1 nm. D_A_ = D_A_ = D_c_ = 10 *µ*m^2^/s. *Sim parameters:* For FPR/NERDSS(Varga et al., 2020), a maximal Δt for each system was limited by the density, values for each crowding fraction: [10^−11^ s, 10^−11^ s, 10^−11^ s, 5×10^−12^ s, 5×10^−12^ s, 10^−12^ s]. N_traj_ =40-80 for each crowding fraction. Another N_traj_ =40-80 were run at another time-step (5 times faster) to verify quantitatively similar results. For eGFRD(Sokolowski et al., 2019), N_traj_ =10 for each crowding fraction. Algorithmic adjustments to standard FPR: First, when an A+B→B+C reaction occurred, the products were kept at their same coordinates as the reactants. Second, at each time-step, particle positions are updated, and they must avoid overlapping (r≥*σ*) all other particles, considering all particles within collision distance (r< *R*_*max*_)(Johnson and Hummer, 2014). Due to the high crowding, we used a more stringent criteria to update particle positions. We defined clusters that contained all the particles with a capacity to possibly overlap all their partners. The positions of all particles in this cluster were simultaneously updated to avoid any overlap. For large clusters, to reduce the number of rejected updates, the time-step was reduced by a factor of 10 and then updated 10 times, to the full Δt.

#### 2. Intermediate Tests

##### 2a Membrane Localization Model

Model Parameters: V=1.1045 *µ*m^3^, A=0.2209 *µ*m^2^, rectangular Cube, 0.47 *µ*m x 0.47 *µ*m surface, height=5*µ*m. [A]_0_=[B]_0_=1 *µ*M or 665 copies, [M]_0_=5.6474*µ*M=17000 /*µ*m^2^, or 3755 copies. [AM]_0_=[BM]_0_=[AB]_0_=[ABM]_0_=. [MAB]_0_=[MABM]_0_=0. Reactions: 1. A+B⇌AB k_on_=0.05 *µ*M^-1^s^-1^. 2. A+M⇌AM k_on_=2 *µ*M^-1^s^-1^. 3. B+M⇌ BM k_on_=2 *µ*M^-1^s^-1^. 4. A+BM ⇌ABM k_on_=0.05 *µ*M^-1^s^-1^. 5. B+MA⇌MAB k_on_=0.05 *µ*M^-1^s^-1^. 6. AB+M ⇌ABM k_on_=2 *µ*M^-1^s^-1^. 7. AB+M⇌MAB k_on_=2 *µ*M^-1^s^-1^. 8. MA+BM⇌MABM 0.0415*µ*m^2^s^-1^. 9. MAB+M⇌MABM 1.6693*µ*m^2^s^-1^. 10. M+ABM ⇌MABM 1.6693*µ*m^2^s^-1^. For all reactions, k_off_=1 s^-1^. At equilibrium, 90.3% of proteins were in complex with one another on the membrane (MABM), or 600 copies, and 1.16% were remaining in solution, or 7.7 copies. *σ*=1 nm for all rxns, D_A_ = D_B_ =50 *µ*m^2^/s. D_AB_ = 25*µ*m^2^/s. D_AM_ = D_BM_=0.495*µ*m^2^/s. D_ABM_ = D_MAB_=0.249*µ*m^2^/s. D_MABM_=0.248*µ*m^2^/s D_M_=0.5 *µ*m^2^/s. All membrane bound had D_z_=0. *Sim parameters:* Single-particle methods Δt=10^−7^s. FPR/NERDSS N_traj_=10. Gillespie N_traj_ =10 Smoldyn N_traj_ =10. We note that the model includes 10 coupled pairwise reactions (three are 3D, four are 3D→2D, and three are 2D), although only 6 equilibrium constants are unique. This is because there are only 3 reactants and multiple thermodynamic cycles that can result in the final assemblies (see e.g. MABM in Fig 5a). To preserve a thermodynamic equilibrium at steady-state, not all rates can be defined independently.

##### 2b Auto Phosphorylation

Model Parameters A: V=0.003 *µ*m^3^. [A]_0_=103 copies, [Ap]_0_=5 copies, [P] _0_=9 copies. [A_Ap]_0_=0, [Ap_P]_0_=0. 1. A→Ap at 21.2 s^-1^. 2. A+Ap→A_Ap at 10^7^M^-1^s^-1^, k_a_=16.7 nm^3^*µ*s^-1^, *σ*=1 nm. 3.A_Ap→Ap+Ap at 200 s^-1^. 4. Ap+P→Ap_P at 8*10^8^M^-1^s^-1^, k_a_=2820 nm^3^*µ*s^-1^, *σ*=1 nm and 5. Ap_P→A+P at 539 s^-1^. D_A_ = D_P_ =100 *µ*m^2^/s. A second spatial stochastic simulation had D_A_ = D_P_ =10 *µ*m^2^/s, which required new bimolecular parameters for rxn 2: k_a_=17.8 nm^3^*µ*s^-1^, *σ*=1 nm, rxn 4: k_a_=11191 nm^3^*µ*s^-1^, *σ*=6 nm. NOTE: the binding radius had to be increased to recover the large macroscopic binding rates for rxn 4. A third spatial stochastic simulation had D_A_ = D_P_ =40 *µ*m^2^/s, which required new bimolecular parameters for rxn 2: k_a_=17.2 nm^3^*µ*s^-1^, *σ*=1 nm, rxn 4: k_a_=3919 nm^3^*µ*s^-1^, *σ*=4 nm.

##### Sim Parameters

Single particle methods: Δt =2*10^−8^s, N_traj_ =2. For D=10, Δt =1*10^−7^s, N_traj_ =2. For D=40, Δt =1*10^−7^s, N_traj_ =2. All simulations were >=200s to obtain statistics on switching times.

B version of model with all rates reduced by 10: 1. 2.12 s^-1^. 2. 10^6^M^-1^s^-1^, k_a_=1.67 nm^3^*µ*s^-1^, *σ*=1 nm. 3. 20 s^-1^. 4. 8*10^7^M^-1^s^-1^, k_a_=282.0 nm^3^*µ*s^-1^, *σ*=1 nm and 5. 53.9 s^-1^. D_A_ = D_P_ =10 *µ*m^2^/s. A second spatial stochastic simulation had D_A_ = D_P_ =100 *µ*m^2^/s, which required for rxn 4: k_a_=140.31 nm^3^*µ*s^-1^, *σ*=1 nm. A third spatial stochastic simulation had D_A_ = D_P_ =1 *µ*m^2^/s, which required for rxn 2: k_a_=1.78 nm^3^*µ*s^-1^, *σ*=1 nm. rxn 4: k_a_=1119.14 nm^3^*µ*s^-1^, *σ*=6 nm.

##### Sim Parameters

Δt =2*10^−7^s, N_traj_ =2. For D=100, Δt =2*10^−8^s, N_traj_ =2. For D=1, Δt =1*10^−6^s, N_traj_ =2. All statistics were collected for at least 2000s, except the fast diffusion constant. With only 100s for D=100, we observed transitions but insufficient statistics to determine trends relative to the other models.

#### 3. Application Tests

##### 3a Circadian Clock Oscillator Model

Model Parameters A: 4.189 *µ*m^3^, cube, or a sphere with R=1 *µ*m. [prmA]_0_=[prmR]_0_=1 copy, or 0.000397 *µ*M. [A]_0_=[R]_0_=[C]_0_= [mRNAA]_0_= [mRNAR]_0_= [prmA.A]_0_=[prmR.A]_0_=0. Reactions: 1. A + R → C k_on_=1204 uM^-1^s^-1^, k_a_=356,263 nm^3^*µ*s^-1^, *σ*=8 nm 2. prmA → prmA + mRNAA k_2_=50 s^-1^. 3. prmA.A → prmA.A + mRNAA k_3_=500 s^-1^. 4. prmR → prmR + mRNAR k_4_=0.01 s^-1^. 5. prmR.A → prmR.A + mRNAR k_5_=50 s^-1^. 6. prmA + A ? prmA.A k_6f_=602 uM^-1^s^-1^, k_6b_=50 s^-1^, k_a_=4889 nm^3^*µ*s^-1^, k_b_=244.5 s^-1^ *σ*=5 nm 7. prmR + A? prmR.A k_7f_=602 uM^-1^s^-1^, k_7b_=100 s^-1^, k_a_=4889 nm^3^*µ*s^-1^, k_b_=489 s^-1^ *σ*=5 nm 8. mRNAA → mRNAA + A k_8_=50 s^-1^. 9. mRNA_R → mRNA_R + R k_9_=5 s^-1^. 10. mRNA_A → NULL k_10_=10 s^-1^. 11. mRNA_R → NULL k_11_=0.5 s^-1^. 12. A → NULL k_12_=1 s^-1^. 13. R→ NULL k_13_=0.2 s^-1^. 14. C→ R k_14_=1 s^-1^. D=10 *µ*m^2^/s (for all 9 species).

B version of the model was run with one modification from the original A model: 13. R→ NULL k_13_=0.05 s^-1^, eliminating oscillations in the deterministic model Fig S3.

C version of this model was created with modifications from original A model: a) promoters were spatially localized to a sphere at the center of the volume of R=0.1 *µ*m. b) D_prmA_=D_prmR_=D_prmA.A_= D_prmA.R_=0. c) [prmA]_0_=[prmR]_0_=1 copy, or here 0.3964 *µ*M. No barrier existed for proteins to access this ‘nucleus’ in the cell center. Full details in SI Methods.

##### Sim parameters

NERDSS Δt=0.5, 2, and 10 *µ*s, and Smoldyn Δt=100 *µ*s. Gillespie N_traj_=10. Smoldyn N_traj_=2, NERDSS N_traj_=2 (600s each) see Table S6 and S7. Time-steps were limited by the fast unbinding reaction of the single promoter, Rxn 7.

##### 3b Model: MinCDE Cell division Model

Model Parameters: Cylindrical volume, h=4 *µ*m, R=0.5 *µ*m. Five species: [MinDT]_0_= 2.1143 *µ*M, or 4000 copies. [MinE]_0_=0.74 *µ*M, or 1400 copies. [MinDD]_0_=[minDt]_0_=[minDt.minE]_0_=0, where lower case indicates membrane bound. Six irreversible reactions. 1. MinDT→minDt, *k*_1_=0.025 um.s^-1^. 2. MinDT + minDt→2minDt, k_2_=0.903s^-1^.uM^-1^. 3.

MinDT+minDt.minE→minDt + minDt.minE, k_3_=0.903 s^-1^.uM^-1^. 4. MinE+minDt→minDt.minE, k_4_=56.0 s^-1^.uM^-1^. 5. minDt.minE→MinDD + MinE, k_5_=0.7s^-1^. 6. MinDD→MinDT, k_6_=1s^-1^. *D*=2.5 *µ*m^2^/s for MinDT, MinDD, and MinE. D=0 *µ*m^2^/s for minDt and minDt.minE in the PDE. Necessary modification for Smoldyn to produce oscillations: *D*=0.05 *µ*m^2^/s for minDt and minDt.minE, initial copies 2000 MinDT, 2000 minDt.

##### Sim parameters

Smoldyn: N_traj_ =4, Δt =2ms.

### Analysis Methods

#### Equilibria and error

For all stochastic methods, we calculated averages and variances of copy numbers in time, over *N* trajectories. We used the standard error of the mean at time points: 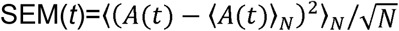. For equilibrium values, we used differences from the mean to determine when equilibrium was reached. We note that for all models, we selected parameters such that the equilibrium was not too close to zero, which is important for assessing clear deviations from the proper equilibrium values.

#### Clock Oscillator timescales

The average time separation between A peaks, between R peaks, and the lag time between A and R peaks was calculated in two ways. First, by finding the peak maxima and simply calculating the separation between neighboring peaks, this was then averaged over all peaks. Second, to calculate time period of peaks, we used a discrete FFT, that was zerσpadded out to 5000 s to increase the resolution of the frequencies sampled. The frequency with the largest coefficient, f_max_, was used to define the period of the oscillation (1/f_max_). The cross-correlation between the A and R time-series was used to identify the lag-time, again keeping the time with the largest correlation coefficient. Both methods produced similar results, and were performed using MATLAB functions. We report in SI tables the periods determined from the find peaks method, with SEMs reporting the calculated 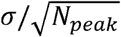.

#### Fit to kinetics of crowding simulations

To define the k_fit_ values for the crowding simulations as a function of packing fraction, we fit the copies of A vs time *A*(*t*) *=A*_0_ exp (*-k*_*fit*_ *B*_*tot*_ *t*), where B_tot_ is 0.008 /nm^3^, and we performed nonlinear fitting to the exponential, rather than the logarithm of A(t). We fit each individual trajectory, and report the mean of these values 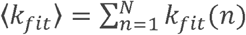, where N is total trajectories. The SEM is calculated based on the variance around this mean value. Fits to the averaged trajectory gave similar values, and fits using the ln(*A*(*t*)) produced the same trends, but with slightly lower *k*_fit_ values. The same approach was used for both FPR and GFRD trajectories.

#### MinCDE timescales

For the MinCDE models, we calculated the time-scales of oscillations by choosing fixed points in space at either end of the cylinder (Fig S7). Once the oscillations were pole-tσpole, we quantified the period of oscillations in time using the same approach as applied to the clock oscillator model above, by finding the peak maxima and taking the average of time separations between adjacent peaks. This produced periods of 43.0±0.7 s for Smoldyn, and 41.3±0.3 s for the PDE.

#### Autophosphorylation State Assignments

To assign stochastic trajectories to states, 1=low phosphorylation, 2=high phosphorylation, we report results based on thresholding the values of A(t) and Ap(t). We created a transition region, where points could be in either 1 or 2 depending on their previous state, to minimize rapid re-crossings between states (see Fig S1). If [A(t)>62+Ap(t)], assign to state 1, elseif [A(t) < 35+Ap(t)] assign state 2, else [if(State(t-1)=*i*, State(t)=*i*], where *i*=[1,2]. We then counted the number of times in each state, length of intervals spent in each state, and transitions between points separated by a time *δ*t to 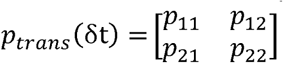. We calculated *p*_*state,i*_ *=n*_*state,i*_ */*(*n*_*state*,1_ *+n*_*state*,2_), and 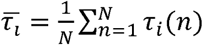, where *N* is the number of intervals spent in state i, each of duration τ_*i*_. Distributions of these residence times are shown in Fig S1. To calculate errors on the probability of each state, each trajectory was split into 10 chunks (length at least the larger residence time), and these 10 state probabilities were used to calculate a mean and standard deviation/SEM.

## ACKNOWLEDGMENTS

This work was funded by the National Institute for Mathematical and Biological Synthesis (NIMBioS). MEJ also gratefully acknowledges computing resources from JHU’s MARCC supercomputer and XSEDE through XRAC MCB150059, and support from an NIH MIRA R35GM133644. **M**.**M**.**S. and T**.**P. acknowledge support by the intramural research program of the National Institute of Allergy and Infectious Diseases, NIAID, NIH**.

