## Supplementary Material for "The Roles of Space and Stochasticity in Computational Simulations of Cellular Biochemistry: Quantitative Analysis and Qualitative Insights"

For

### Supporting Methods

**A. Derivation for autophosphorylation model rate balance analysis** shown in Figure 6b. The production and degradation terms for Ap can be derived from its rate equation:

$$\frac{d\ Ap}{dt} = \underbrace{k_1 \cdot A + k_3 \cdot AAp}_{\text{production}} - \underbrace{k_4 \cdot Ap \cdot P}_{\text{degradation}}$$

where *Ap* is the phosphorylated kinase, *A* is the unphosphorylated kinase, *P* is the phosphatase, and *AAp* is a dimer of kinases. To solve for the production and degradation terms each as a function of *Ap* only, we use the conservation of total kinase $K_T$ and phosphatase $P_T$:

$$P + A_pP = P_T$$

$$A + Ap + 2AAp + ApP = K_T$$

Along with the rate equations for *ApP*, the dimer of kinase and phosphatase, and *AAp*, the kinase dimer, solved for at steady-state:

$$\frac{d\ ApP}{dt} = k_4 \cdot P \cdot Ap - k_5 \cdot ApP = 0$$

$$\frac{d\ AAp}{dt} = k_2 \cdot A \cdot Ap - k_3 \cdot AAp = 0$$

Using these 5 equations, we recover:

$$production = \frac{(k_1 + k_2Ap)\left(K_T - Ap - \frac{P_T \cdot Ap}{{k_5}/{k_4} + Ap}\right)}{1 + {2k_2}/{k_3}\ Ap}$$

$$degradation = \frac{k_5 \cdot Ap \cdot P_T}{{k_2}/{k_3} + Ap}$$

For particular parameter sets, the fixed points are then determined for values of *Ap* where the production is equal to the degradation.

**B. Additional Model description details for spatial clock model simulations with localized promoters.**
The reaction network is unchanged. For this model, promoters are localized to the center of the system, with $D_{PRMA} = D_{PRMR} = 0\ \mu m^2/s$ (bound and unbound).
V1. $D_A = D_R = D_{RNA} = D_{RNP} = 10\ \mu m^2/s$. $D_{A.R} = 10\ \mu m^2/s$.
V2. $D_A = 5\ \mu m^2/s$. $D_R = 50\ \mu m^2/s$. $D_{RNA} = D_{RNP} = 5\ \mu m^2/s$. $D_{A.R} = 4.5\ \mu m^2/s$.
V3. $D_A = 2\ \mu m^2/s$. $D_R = 50\ \mu m^2/s$. $D_{RNA} = D_{RNP} = 5\ \mu m^2/s$. $D_{A.R} = 1.9\ \mu m^2/s$.
For the NERDSS simulations, note the binding radii had to increase to accommodate the slowing diffusion.
V1. A+R: $k_a = 356262.9\ nm^3 \mu s^{-1}$, $\sigma = 8$ nm. A+PrmA: $k_a = 189336\ nm^3 \mu s^{-1}$, $\sigma = 8$ nm, $k_b = 9466.8\ s^{-1}$ A+PrmR: $k_a = 189336\ nm^3 \mu s^{-1}$, $\sigma = 8$ nm, $k_b = 18933.6\ s^{-1}$
V2. A+R: $k_a = 4747.72\ nm^3 \mu s^{-1}$, $\sigma = 5$ nm. A+PrmA: $k_a = 189336\ nm^3 \mu s^{-1}$, $\sigma = 16$ nm, $k_b = 9466.8\ s^{-1}$ A+PrmR: $k_a = 189336\ nm^3 \mu s^{-1}$, $\sigma = 16$ nm, $k_b = 18933.6\ s^{-1}$
V3. A+R: $k_a = 5156.4\ nm^3 \mu s^{-1}$, $\sigma = 5$ nm. A+PrmA: $k_a = 189336\ nm^3 \mu s^{-1}$, $\sigma = 40$ nm, $k_b = 9466.8\ s^{-1}$ A+PrmR: $k_a = 189336\ nm^3 \mu s^{-1}$, $\sigma = 40$ nm, $k_b = 18933.6\ s^{-1}$

*Sim Parameters: V1:* $\Delta t = 2*10^{-6}$s. V2: $\Delta t = 0.5$ and $2*10^{-6}$s. V3: $\Delta t = 0.5$ and $2*10^{-6}$s.

### Supporting Figures:

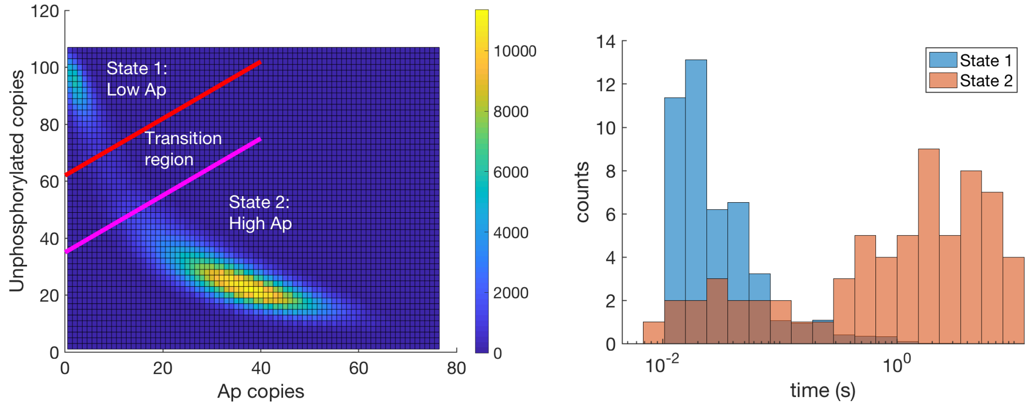


**Fig S1. State assignment and analysis of autophosphorylation model.** a) 2D histogram of copy numbers of both of Ap and A (unphosphorylated). State assignments were determined from the 2D vector of copy numbers at each time point. To minimize rapid crossings across a single threshold line, which biased towards very short residence times, we created two thresholds and a transition region. All values of A above the red line are in State 1, and all below the pink line are in State 2. Points in the transition region can be in either state, depending on where they originated. Points coming from State 1 remain in State 1 until they cross the pink link. Points coming from State 2 remain in State 2 until they cross the Red line. This distribution is representative of most trajectories, with ~88% of points assigned to State 2. b) Histogram of counts of residence times spent in either state. Bin widths are even on a log10 scale. For both plots, data is from a Gillespie simulation for 200s, with points saved every $10^{-4}$s. The analysis above resulted in 68 transitions out of each state.

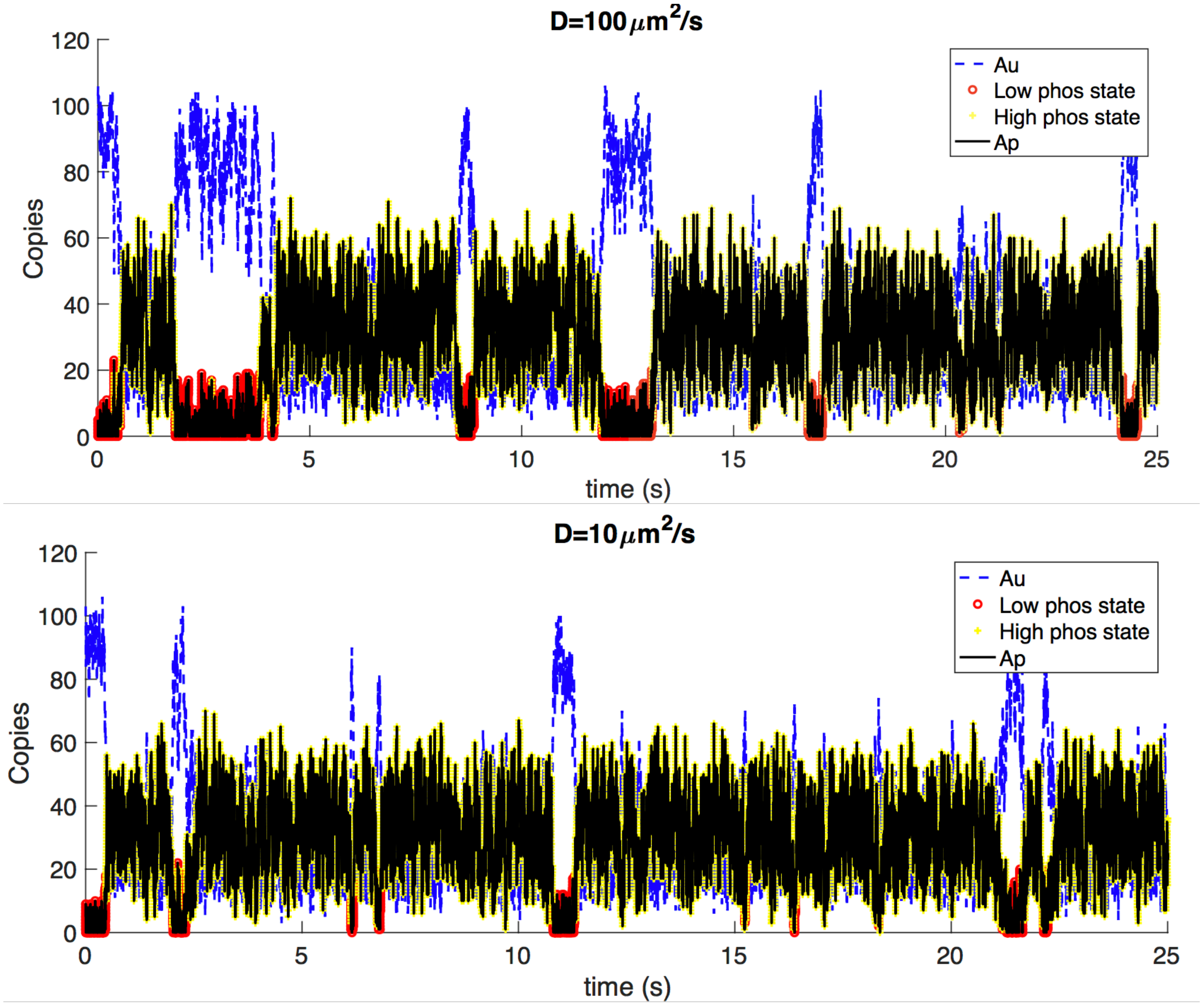


**Fig S2. Autophosphorylation model shows switching in stochastic single-particle simulations with distinct diffusion coefficients.** (Top) From NERDSS simulations with D=100 $\mu m^2$/s, the kinase is in the low phosphorylation state 17% of the time (red pluses), in intervals of 0.3±0.04s. The high state (yellow circles) for 1.46±0.17s intervals. The statistics were collected from two 100s trajectories. (Bottom) For D=10 $\mu m^2$/s, the results are similar, but the kinase spends 9% of time in the low state, in intervals of 0.18±0.02s, and remainder in the high state with intervals of 1.73±0.2s. The statistics were collected from a 200s trajectory, with similar results from independent trajectories. In both plots, black data is the copies of phosphorylated kinase, and blue data is copies of the unphosphorylated kinase. State assignments were made based on thresholding across two transition lines (Methods and Fig S1).

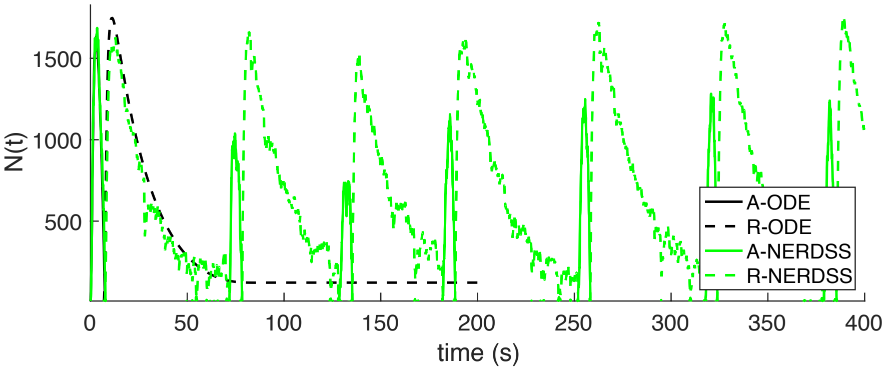


**Fig S3. Clock oscillator model with slower decay of repressor protein.** Compared with Fig. 3, here, the decay of the repressor is slowed from, 0.2 to 0.05$s^{-1}$, causing oscillations to disappear in the deterministic simulation (black curves) but persist in the stochastic single-particle simulations (green curves), as observed in the original publication[1]. The single-particle NERDSS simulations[2] were run for 1000s, producing oscillations with periods of 63±2.8 and 63±2.8s (SEM over 16 peaks), or more than twice as slow as the original model (25s). The lag between A and R slowed from 6s to 7.6s±0.3.

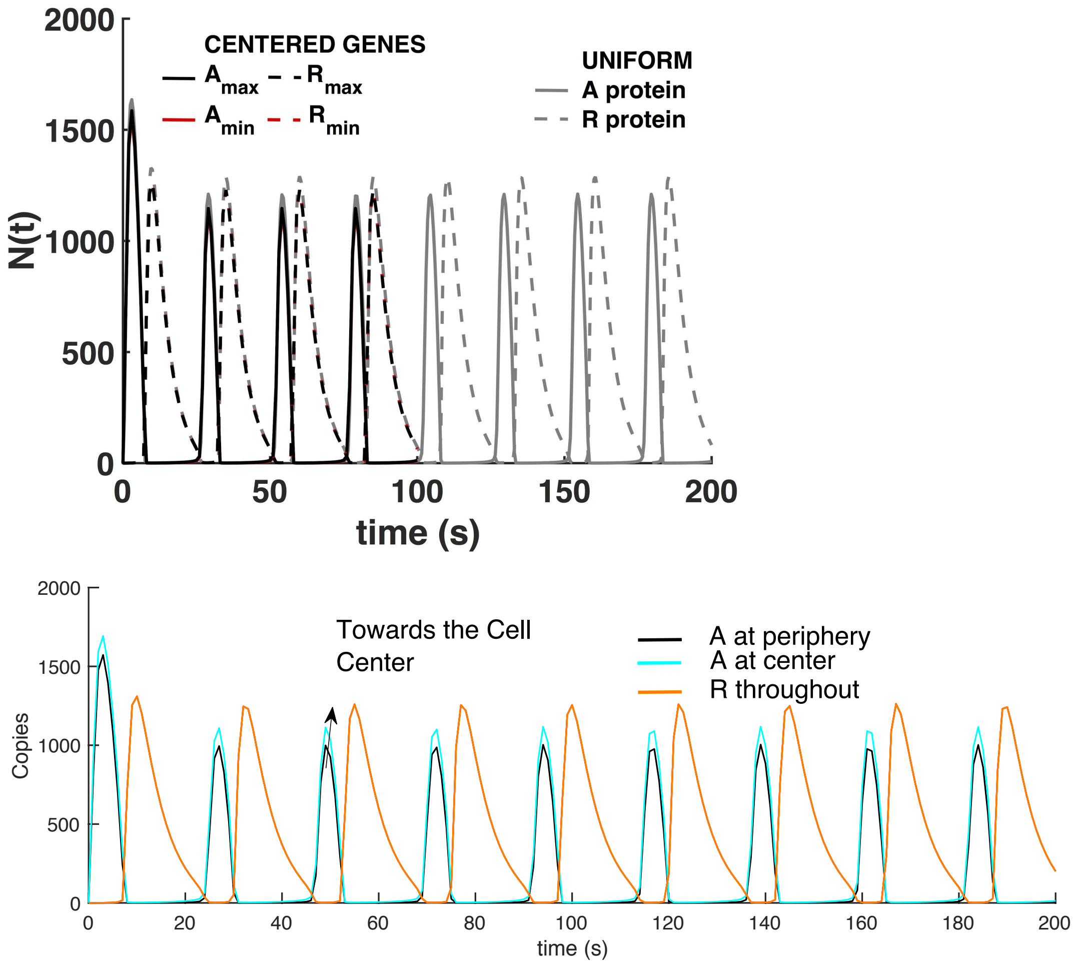


**Fig S4. Clock Oscillator model with localized genes.** Localizing and immobilizing the DNA (prmR and prmA) to the center of the volume in PDE models. Top: this does not affect the oscillations produced in a

PDE solution (black curves) for the small cell size studied here (R=1$\mu$m), when $D_A$=10 $\mu m^2$/s.  Across the width of the cell, the density of molecules is nearly uniform, showing undetectable differences between the maximum and minimum values of A or R reached for each time point. For reference, we plot the ODE solution that thus has no spatial dependence (gray curves). Bottom: With slowing A, where $D_A$=2 $\mu m^2$/s, we do start to see increased localization of A in the cell center near the promoter (cyan vs black), causing a shortening of the oscillation time from 25s to 22.6s.

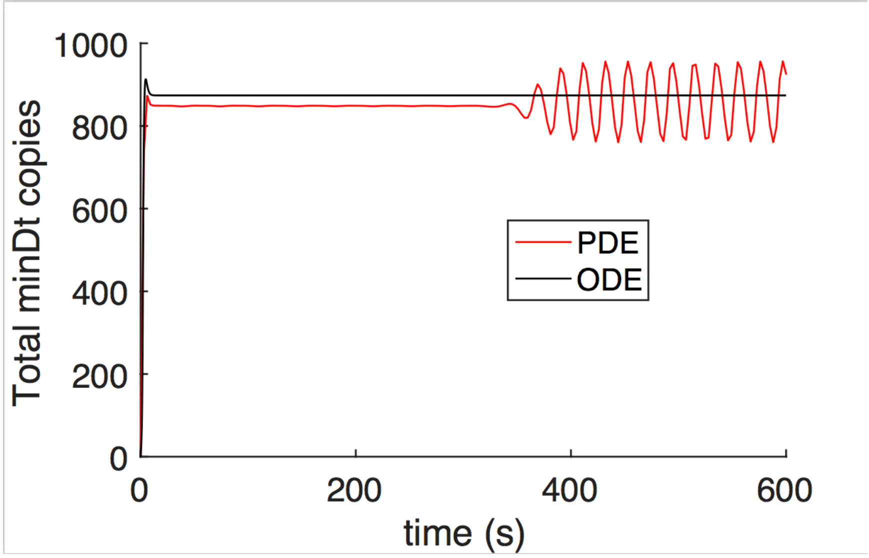


**Fig S5. For MinCDE model, no MinD-ATP$^{2D}$ oscillations occur in nonspatial model.** In the ODE model, no spatial oscillations are possible for MinD-ATP$^{2D}$, but no temporal oscillations occur either. For the PDE, the total MinD-ATP$^{2D}$ copies are constant whilst their spatial oscillations are symmetric, but their total copies also oscillate when spatial oscillations are pole-to-pole.

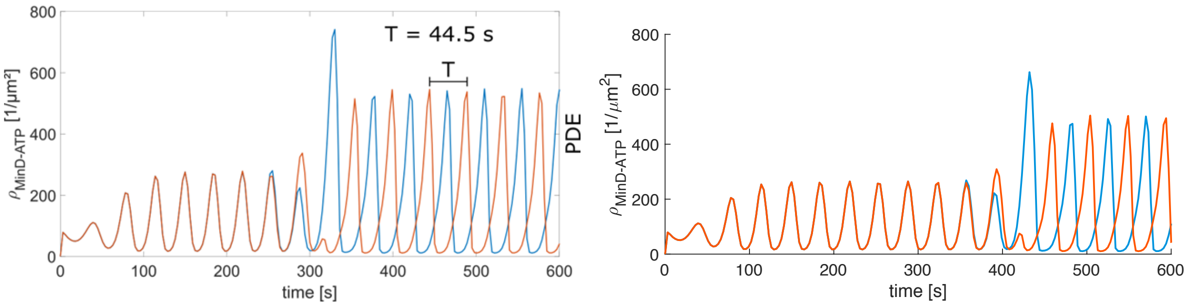


**Fig S6. MinCDE model dependence on initial copies and error tolerance.** a) By increasing the initial copy numbers of [MinD-ATP, MinE] from [2.1143,0.74] to [2.5, 0.875] $\mu$M, the symmetric oscillations at the beginning of the simulation are much more apparent relative to Fig S7 and Fig 8 main text. We note the presence of oscillations are also sensitive to the ratio of MinD-ATP/MinE, and a sufficient total concentration of both (e.g. no oscillations with half of the original copies).   b) The species are initially uniform in the solution volume and oscillations are symmetric (blue and orange copies on exact opposite points of cylinder). Symmetry breaking occurs due to numerical precision errors. On the right, we increased the [absolute, relative error tolerances] of the PDE from [1e-9, 1e-7] to [1e-10, 1e-9], causing the symmetric oscillations to persist out to almost ~400s, instead of only ~300s, showing that the pole-to-pole oscillations are clearly more stable and robust. Increasing the mesh size by a factor of two also delays the symmetry breaking to ~320s.

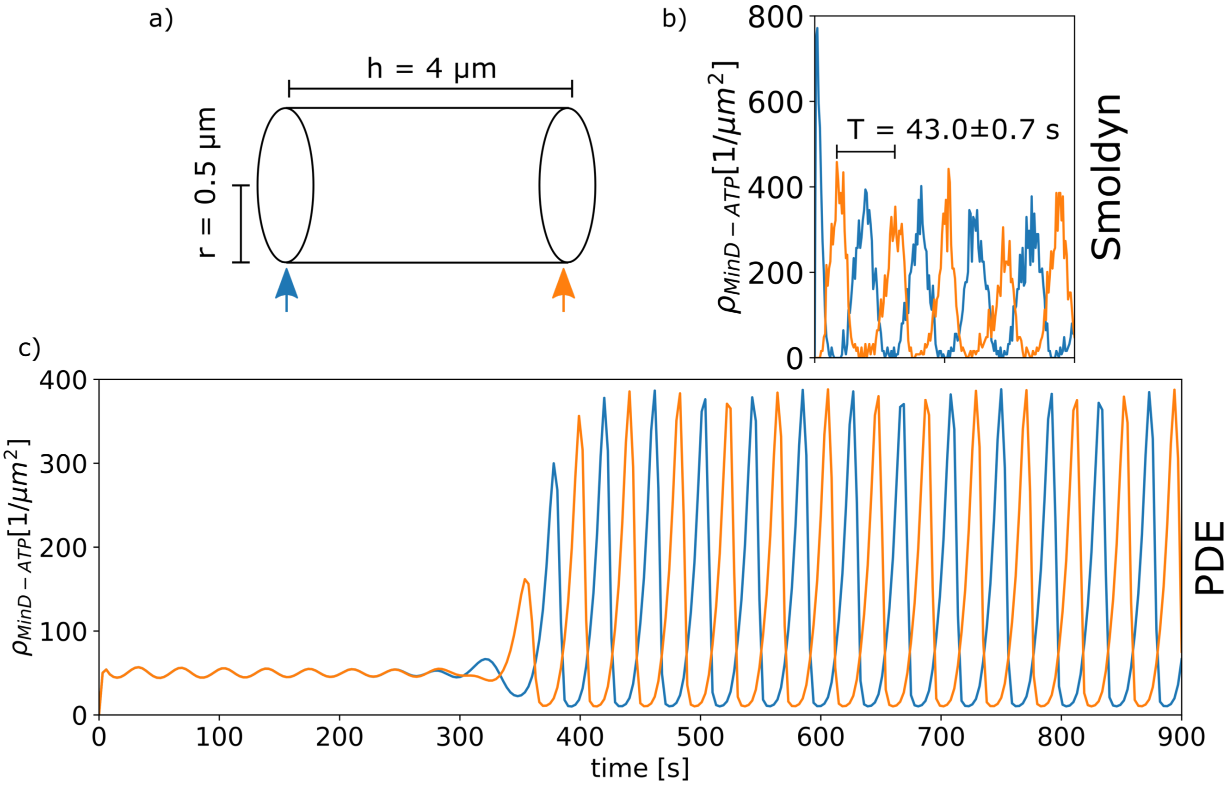


**Fig. S7 MinCDE model, time-dependent minDt oscillations at cell ends**. (a) sketch of the cell with particles at both poles being tracked, same model as Fig 4. (b) Pole-to-pole oscillation in the Smoldyn model. The high peak at t = 0 s shows the asymmetric initialization. (c) Symmetric oscillation of the PDE model up to 250 s. Afterwards the oscillation switches to a pole-to-pole mode. The period of these oscillation is roughly the same in the Smoldyn (43.0 ± 0.7 s) and PDE (41.3 ± 0.3 s) model.

### Supporting Tables

#### Table S1. Categorized software tools

| TOOL/Method | Rate eqs:ODE | CME/Gillespie | RD: PDE | RDME | Single-particle |
|---|---|---|---|---|---|
| Virtual Cell | ■ | ■ | ■ | | ■ |
| ML-Space | | ■ | | ■ | ■ |
| PSpatioCyte | | ■ | | ■ | ■ |
| Lattice Microbes | | | | ■ | |
| stochSS/RDME | | | | ■ | |
| Simmune | | | ■ | | |
| MCell | | | | | ■ |
| NERDSS | | | | | ■ |
| Smoldyn | | | | | ■ |
| EGFRD | | | | | ■ |

#### Table S2. Software tools and features

| | | Single particle. | | Species features | | | | Boundary features | | | | | User-friendliness | |
|---|---|---|---|---|---|---|---|---|---|---|---|---|---|---|
| | | Smoluchowski-based | Single-particle Kinetic Monte Carlo | Excluded Volume | Intermolecular force | Multi-site molecules | Orientation-dependence | Curved membranes | Realistic Cell Morphology | Compartments | Movable boundaries | Membrane mechanics | Rule-Based capacity or multi-valency | GUI set-up, run, analyse |
| PDE | VCell/PDE solver | – | – | – | – | – | – | ■ | ■ | ■ | ■ | | ■ | ■ |
| PDE | Simmune | – | – | – | – | – | – | ■ | ■ | ■ | ■ | | ■ | ■ |
| RDME | stochSS/RDME | – | – | – | – | – | – | ■ | ■ | ■ | ■ | | ■ | ■ |
| RDME | Lattice Microbes | – | – | – | – | – | – | | ■ | ■ | | | ■ | ■ |
| Single-particle | pSpatioCyte | see 1 | | ■ | | | | ■ | ■ | ■ | | | ■ | |
| Single-particle | Smoldyn | see 2 | | see 3 | | | | ■ | ■ | ■ | | | ■ | see 4 |
| Single-particle | MCell | | ■ | | | | | ■ | ■ | ■ | | | ■ | ■ |
| Single-particle | VCell/Spring Salad | | ■ | ■ | | ■ | | | | | | | ■ | ■ |
| Single-particle | Readdy 2 | | ■ | ■ | ■ | ■ | | ■ | ■ | | | | ■ | ■ |
| Single-particle | NERDSS | ■ | | ■ | ■ see 3 | ■ | ■ | ■ | | | | | ■ | ■ |
| Single-particle | eGFRD | ■ | | ■ | | | | | | | | | ■ | |

**KEY:** DARK shade=true. GRAY=not-applicable. **1.** On lattice. **2.** Approximates Smoluchowski model. **3.** Not standard. **4.** In VCell

**Table S3. Wall time for simulation tools applied to the clock model**

| | **Total Wall time for 30s Clock model sim. (s)** | **Wall time per iteration (s)** | **Time-step (s)** |
|---|---|---|---|
| **ODE** | 2 | - | - |
| **Gillespie** | 2 | - | - |
| **PDE** | 53 | - | - |
| **Smoldyn** | 166246 | 0.0055 | $10^{-6}$ |
| **NERDSS** | 11422 | 0.0038 | $10^{-5}$ |
| **Mcell** | 12179 | 0.0004 | $10^{-6}$ |

**Table S4. Autophosphorylation Model State properties**

| | $P_{state\ 1}$ | $P_{state\ 2}$ | $\tau_1$ (s) | $\tau_2$ (s) | $N_{transition}$ |
|---|---|---|---|---|---|
| SSA (200s) | 0.12±0.02$^{+}$ | 0.88±0.02$^{+}$ | 0.34±0.06$^{\blacktriangle}$ | 2.6±0.36$^{\blacktriangle}$ | 68 |
| NERDSS (D100) (2x100s) | 0.17±0.026$^{+}$ | 0.83±0.026$^{+}$ | 0.3±0.04$^{\blacktriangle}$ | 1.46±0.17$^{\blacktriangle}$ | 113 |
| NERDSS (D40) | 0.09±0.02$^{+}$ | 0.91±0.02$^{+}$ | 0.2±0.025$^{\blacktriangle}$ | 1.93±0.25$^{\blacktriangle}$ | 94 |
| NERDSS (D10) (200s) | 0.09±0.02$^{+}$ | 0.91±0.02$^{+}$ | 0.18±0.02$^{\blacktriangle}$ | 1.73±0.2$^{\blacktriangle}$ | 105 |

$^{+}$SEM values reported over 10-20 chunks of the full trajectory. $^{\blacktriangle}$SEM values reported as $\sigma/\sqrt{N_{transition}}$.

**Table S5. Autophosphorylation Model B State properties, with all rates reduced by 10**

| | $P_{state\ 1}$ | $P_{state\ 2}$ | $\tau_1$ (s) | $\tau_2$ (s) | $N_{transition}$ |
|---|---|---|---|---|---|
| SSA (2000s) | 0.13±0.025$^{+}$ | 0.87±0.025$^{+}$ | 3.6±0.66$^{\blacktriangle}$ | 25.7±3$^{\blacktriangle}$ | 69 |
| NERDSS (D100) (**103s: N.S.**) | 0.27±**0.2**$^{+}$ | 0.73±**0.2**$^{+}$ | 4±**2.8**$^{\blacktriangle}$ | 15±**11.0**$^{\blacktriangle}$ | **6** |
| NERDSS (D10) (2x1000s) | 0.18±0.02$^{+}$ | 0.82±0.02$^{+}$ | 2.7±0.3$^{\blacktriangle}$ | 12.3±1.2$^{\blacktriangle}$ | 149 |
| NERDSS (D1) (5180s) | 0.07±0.008$^{+}$ | 0.93±0.008$^{+}$ | 1.4±0.1$^{\blacktriangle}$ | 19±1.4$^{\blacktriangle}$ | 253 |

$^{+}$SEM values reported over 10 chunks of the full trajectory. $^{\blacktriangle}$SEM values reported as $\sigma/\sqrt{N_{transition}}$. N.S. Not significant—trajectories are not long enough.

**Table S6. Clock model time-scales where *D*=10$\mu$m$^2$/s for all species.**

| | A period (s) | R period (s) | Lag-time (s) | Sim. Time (s) | $N_{peaks}$ |
|---|---|---|---|---|---|
| ODE | 25.2±0.1$^{+}$ | 25.1±0.1$^{+}$ | 6±0.1$^{+}$ | 200 | 8 |
| PDE | 25.3±0.2$^{+}$ | 25±0$^{+}$ | 5.9±0.2$^{+}$ | 200 | 8 |
| Gillespie | 25±0.4$^{+}$ | 25±0.4$^{+}$ | 6±0.1$^{+}$ | 200x10 traj. | 74 |
| NERDSS | 24.4±0.4$^{+}$ | 24.3±0.4$^{+}$ | 6.1±0.1$^{+}$ | 600 x2 traj. | 49 |
| Smoldyn | 25.9±0.9$^{+}$ | 25.7±0.9$^{+}$ | 5.7±0.2$^{+}$ | 200x2 traj. | 16 |

$^{+}$SEM values reported over $N_{peaks}$.

**Table S7. Clock model time-scales for localized and immobile promoters.**

| | A period (s) | R period (s) | Lag-time (s) | Sim. Time (s) | $N_{peaks}$ |
|---|---|---|---|---|---|
| PDE ($D_A$=10) | 25.3±0.2$^{+}$ | 25±0$^{+}$ | 5.9±0.2$^{+}$ | 200 | 8 |
| PDE ($D_A$=5) | 24.5±0.3$^{+}$ | 24.3±0.1$^{+}$ | 5.9±0.2$^{+}$ | 200 | 9 |
| PDE ($D_A$=2) | 22.6±0.3$^{+}$ | 22.5±0.1$^{+}$ | 5.8±0.2$^{+}$ | 200 | 10 |
| NERDSS ($D_A$=10) | 24.7±0.7$^{+}$ | 24.7±0.7$^{+}$ | 6.1±0.1$^{+}$ | 1000 $\Delta$t=10$\mu$s | 41 |
| NERDSS ($D_A$=10) | 24.5±1.2$^{+}$ | 24.4±1.9$^{+}$ | 5.7±0.5$^{+}$ | 200 $\Delta$t =2$\mu$s | 9 |
| NERDSS ($D_A$=5) | 23.7±0.8$^{+}$ | 23.5±0.8$^{+}$ | 5.7±0.2$^{+}$ | 260 $\Delta$t =0.5 $\mu$s | 12 |
| NERDSS ($D_A$=5) | 23.8±0.5$^{+}$ | 23.8±0.6$^{+}$ | 5.8±0.2$^{+}$ | 830 $\Delta$t =2 $\mu$s | 34 |
| NERDSS ($D_A$=2) | 22.3±0.8$^{+}$ | 22.5±1$^{+}$ | 5.9±0.3$^{+}$ | 250 $\Delta$t =0.5 $\mu$s | 11 |
| NERDSS ($D_A$=2) | 23±0.4$^{+}$ | 23±0.5$^{+}$ | 5.7±0.2$^{+}$ | 880 $\Delta$t =2 $\mu$s | 37 |

$^{+}$SEM values reported over $N_{peaks}$. Oscillation periods were the same at the center and periphery of the cell in the PDE. NERDSS periods are from spatially averaged copies vs time.
